# Outer Membrane Vesicles Secreted by *Bacteroides fragilis* Inhibit CFTR Chloride Secretion by Human Colon Organoids

**DOI:** 10.1101/2025.08.19.671135

**Authors:** Roxanna Barnaby, Amanda Nymon, Paige Salerno, Carolyn Roche, Young Ah Goo, Byoung-Kyu Cho, Timothy B. Gardner, Zdenek Svindrych, Douglas J. Taatjes, Thomas H. Hampton, Benjamin Ross, Bruce A. Stanton

## Abstract

The goals of this study were to develop a model to study host pathogen interactions in primary human colon organoids and to test the hypothesis that *Bacteroides fragilis* toxin (BFT-2) secreted in outer membrane vesicles (OMVs) modulates mucosal immunity and CFTR Cl^-^ secretion. Since Bacteroides species reside in mucus, OMVs are likely to represent a mechanism of communication between Bacteroides and the host. Two strains of Bacteroides were studied, Enterotoxigenic *Bacteroides fragilis* (ETBF), which produces BFT-2, and the non-toxigenic *Bacteroides fragilis* strain NCTC 9343 (NTBF) that does not produce BFT-2. We also utilized two additional strains of *Bacteroides fragilis:* one in which *bft*-2 was knocked out (ETBF Δ*bft*), and one that was engineered to contain *bft-2* (NTBF+*bft)*. We report that *Bacteroides fragilis* OMVs reduced CFTR Cl*^-^* secretion but had no effect on tight junction or cell adhesion proteins, transepithelial resistance (TER) or cytokine secretion by primary human colon organoids. NTBF OMVs containing BFT-2 were more effective in reducing CFTR Cl*^-^* secretion than NTBF lacking BFT-2. We conclude that OMVs secreted by Bacteroides can be an important mechanism of host pathogen interactions in the colon by reducing CFTR Cl^-^ secretion.

## Introduction

The human gastrointestinal tract is colonized by ∼100 trillion bacteria, including several members of the genus Bacteroides, which comprise approximately 30% of the microbiota ^1–5^. Although *Bacteroides fragilis* (*B. fragilis*) is one of the least abundant species, Enterotoxigenic *B. fragilis* (ETBF) is a pathogenic bacterium that induces diarrhea, colitis and causes tumor formation through the secretion of Bacteroides fragilis toxin (BFT-2), a metalloprotease ^3,6–8^. Many, but not all, studies on colon cell lines have shown that recombinant BFT-2 rapidly (∼1-3 hrs.) causes cell damage and disrupts cell adhesion by cleaving E-cadherin, a cell adhesion protein, leading to an increase in epithelial paracellular permeability, IL-8 secretion ^3,9–11,12^, fluid secretion driven by CFTR Cl^-^ channels ^13,14^ and diarrhea ^15–17^. However, in some studies on colon cell lines, E-cadherin remained unchanged after exposure to ETBF supernatant ^9^. The effect of NTBF and ETBF OMVs on IL-8 secretion and E-cadherin in colon cell lines is dose, strain and time dependent ^9,12,18^. It has been suggested that the BFT-2-induced increase in permeability leads to paracellular movement of bacteria and bacterial products from the lumen of the colon to the submucosal layer and blood, an effect that can train the immune system ^2^. Recombinant BFT-2 applied to the apical side of HT 29/C1, HT-29, T84, CMT93 or MDCK cell lines decreases transepithelial resistance (TER) and cleaves E-cadherin ^9,11,15,17,19,20^. One study on the T84 colon cell line reported no effect of BFT-2 on short circuit current but did report that BFT-2 reduced TER ^20^.

Many Gram-negative bacteria that reside in the mucus lining of the colon secrete outer membrane vesicles (OMVs) ^12,21–26^ that stimulate inflammation and decrease ion transport, including CFTR Cl^-^ secretion by epithelial cell lines ^21–23,27^. Therefore, it is likely that communication between *B. fragilis* and human colon cells is also mediated by the secretion of OMVs and other soluble factors including MUC-2, antimicrobial peptides, and anti-inflammatory cytokines ^7,28,29^. However, nothing is known about the effects of BFT-2-containing OMVs secreted by *Bacteroides* on primary human colon organoids.

In contrast to ETBF, the Nontoxigenic *B. fragilis* (NTBF) strain does not express *bft-2* and it promotes gut homeostasis through the production of short-chain fatty acids and polysaccharide A, stimulates barrier-protective molecules including ZO-1 and MUC-2, as well as the secretion of antimicrobial peptides, and anti-inflammatory cytokines ^1,7,30^.

A recent review questioned the relevance of cell lines as models to elucidate the physiology and pathophysiology of the colon since cell lines are rarely normal, and contain several mutations and chromosomal defects that enable their persistence ^31^. In particular, it has been reported that the gene expression pattern of the Caco-2 cell line, which is studied as a surrogate of the colon, was similar to that of epithelial cells of the small intestine ^32^. Since, as noted above, colon cell lines do not produce consistent responses to recombinant BFT-2 ^3,9,10^, one of the most important challenges involves the assessment of BFT-2 secreted in OMVs on primary human colon organoids ^1^. To elucidate the biological activity of BFT-2 secreted in OMVs, we obtained colon biopsies from multiple healthy human donors to develop differentiated two-dimensional monolayers of primary colon organoids. Primary colon organoids contain the multiple cell populations found in native tissue, compared to cell lines, which do not fully differentiate ^31^. Accordingly, the goals of this study were to develop a model to study host pathogen interactions in primary human colon organoids, and to test the hypothesis that BFT-2 secreted in OMVs modulates mucosal immunity, TER and CFTR Cl^-^ secretion, a major driver of fluid secretion in the colon ^14^. We report that OMVs had no effect on cell adhesion or tight junction proteins, notably E-cadherin and claudin-2, TER, or sodium reabsorption, but did inhibit forskolin-stimulated CFTR Cl^-^ secretion, an effect predicted to decrease fluid secretion by the colon. NTBF OMVs containing BFT-2 had a more inhibitory effect on CFTR Cl^-^ secretion by primary human colonoids than NTBF OMVs lacking BFT-2. In addition, OMVs were not pro-inflammatory as determined by measuring the secretion of 48 cytokines. OMVs also reduced CFTR Cl^-^ secretion by T84 cells, a colon cell line, but did not alter TER. These results demonstrate that OMVs and BFT-2 secreted by *B. fragilis* have very different biological effects on primary human colon organoids compared to colon cell lines exposed to recombinant BFT-2.

## Results

### Isolation and characterization of OMVs secreted by Bacteroides

To test the hypothesis that BFT-2 in OMVs leads to a disruption in the integrity of epithelial barriers and modulates CFTR Cl^-^ secretion, we utilized four strains of *B. fragilis*, including ETBF and ETBF in which the *bft-2* gene was deleted (ETBF Δ*bft*), NTBF, a strain that does not express *bft-2,* and NTBF in which the *bft*-2 gene was expressed (NTBF+*bf*t). Nanoparticle tracking analysis (NTA), which measures the size of vesicles by dynamic light scattering, revealed that the size of OMVs isolated from bacterial cultures was similar for most strains, except that ETBF OMVs were significantly smaller than NTBF OMVs (**Figure 1**).

**Figure 1:**
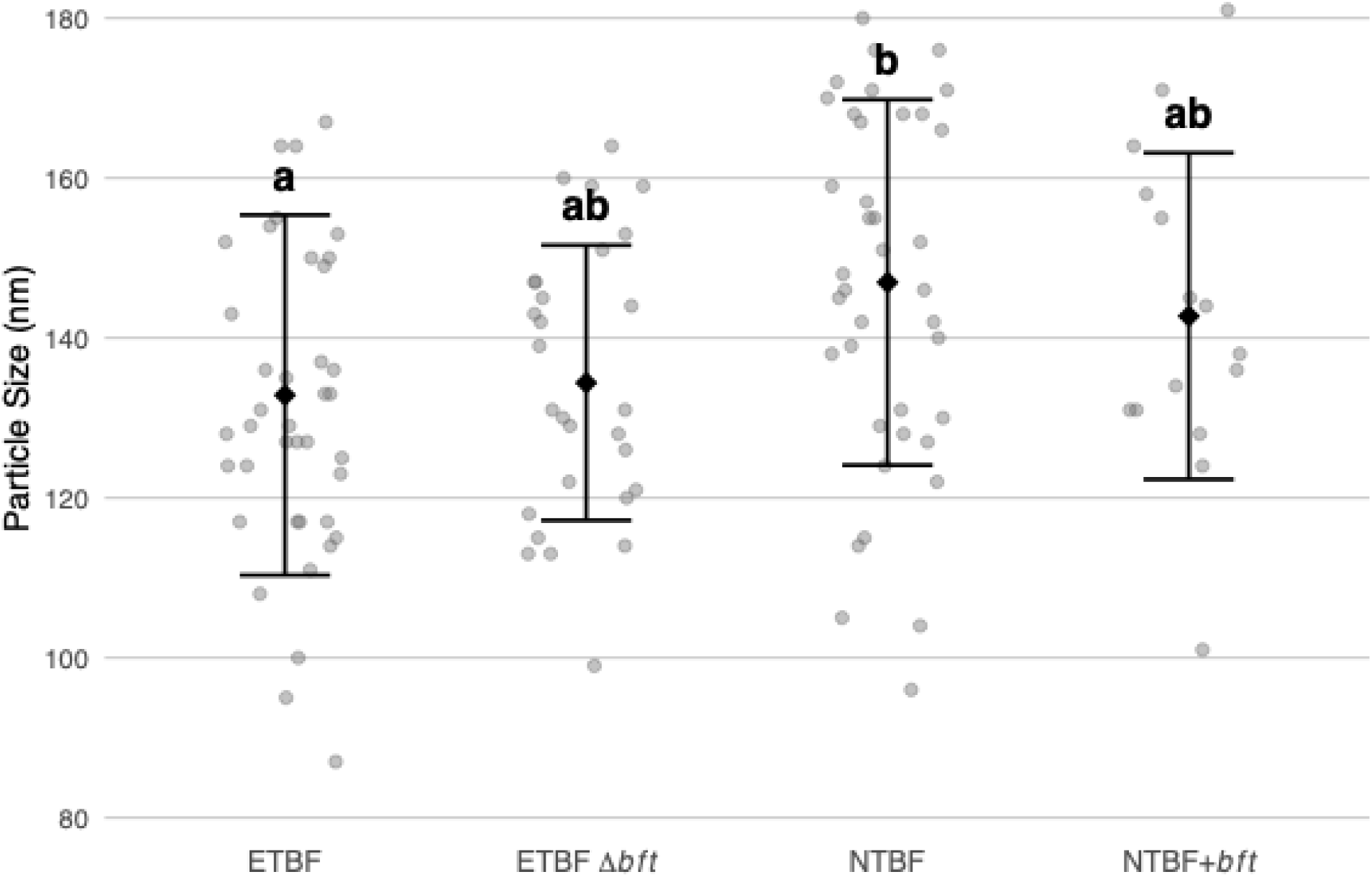
Bacteroides OMV particle size determined by NTA. Individual measurements are shown as gray circles, with group means (black diamonds) and error bars (±SD). Treatments: ETBF, ETBF Δ*bft*, NTBF, and NTBF+*bft*. Letters above bars indicate statistical groupings based on one-way ANOVA with Tukey’s HSD post-hoc test; groups sharing the same letter are not significantly different (P > 0.05), while groups with different letters are significantly different (P < 0.05).

According to NTA analysis, the number of OMVs in each bacterial culture was: 1.4 x 10^11^ particles/ml for ETBF, 2.8 x 10^11^ particles/ml for ETBF Δ*bft,* 2.0 x 10^11^ particles/ml for NTBF and 2.2 x 10^11^ particles/ml for NTBF+*bft*. To confirm that we isolated OMVs, transmission electron microscopy (TEM) was utilized to image OMVs (**Figure 2**). OMVs were not observed in process control (PC, bacterial culture media not exposed to bacteria and run through the OMV isolation process); however, OMVs were observed in media isolated from ETBF, NTBF, ETBF Δ*bft* and NTBF+*bft* (**Figure 2**). The mean diameter (±SEM) of OMVs as measured by TEM were similar in all groups (P>0.77): ETBF [(59.0 nm ± 2.0 nm, (n=31)], ETBF Δ*bft* [(63.2 ± 3.6 nm (n=21)], NTBF [(60.2 ± 4.0 nm, (n=14)] and NTBF+*bft* [(58.0 ± 5.0 nm (n=11)]. Most Bacteroides OMVs had typical morphology, but we also observed some collapsed vesicular forms and the presence of some irregularly shaped vesicles, similar to images published recently for Bacteroides OMVs secreted by ETBF and NTBF ^33^. As expected, the size of OMVs was assay dependent: dynamic light scattering tends to overestimate size, whereas TEM tends to underestimate size ^34,35^.

**Figure 2.**
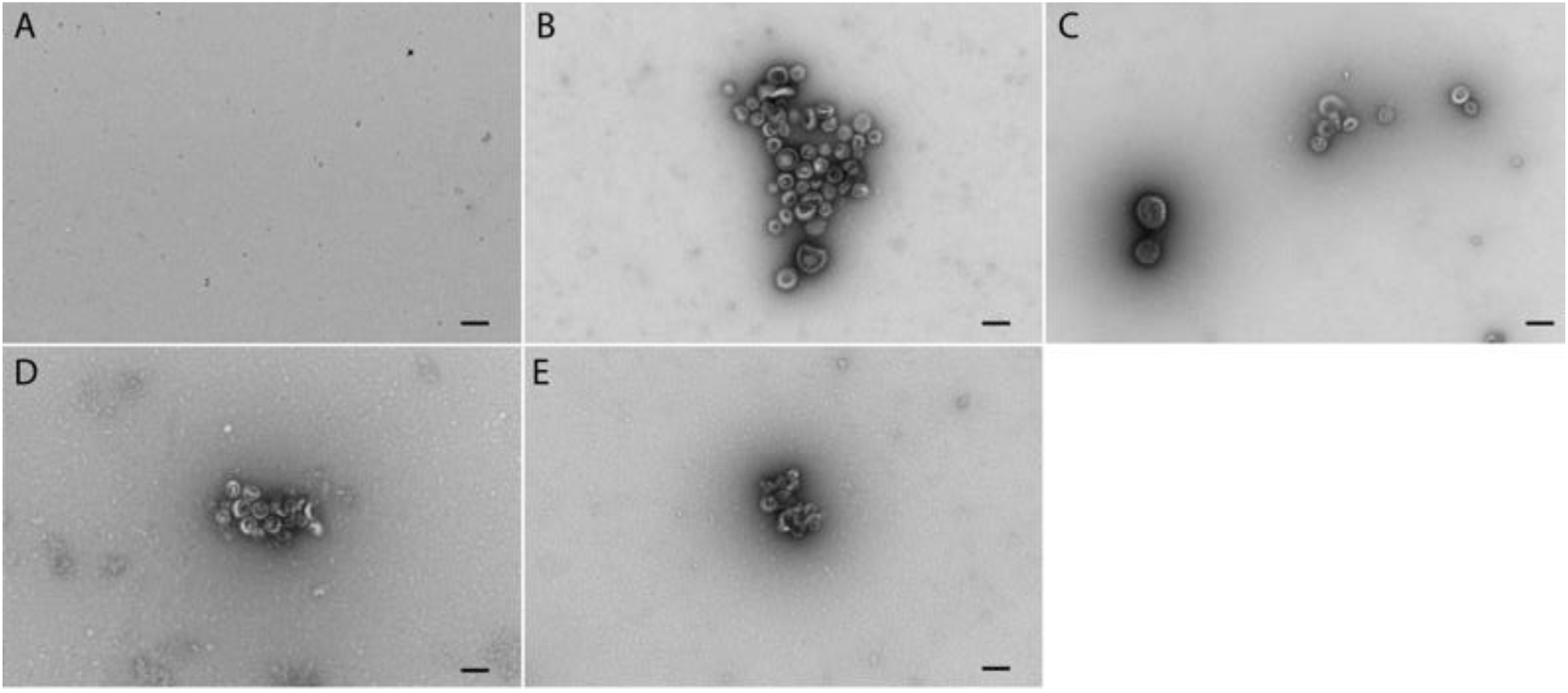
Representative transmission electron microscopy images of negatively stained Bacteroides OMVs. (A) Process control. (B) ETBF. (C) ETBF Δ*bft*. D. NTBF. E. NTBF+*bft*. Images taken by an observed blinded to the origin of the OMVs. Scale bars 100 nm.

Analysis of the proteome of OMVs was conducted to confirm that BFT-2 was present in ETBF and NTBF+*bft* and absent in ETBF Δ*bft* and NTBF. Results of the proteomics analysis of OMVs are presented in **Figure 3 and** Supplemental Tables 1-4. We detected 568 proteins in NTBF OMVs and 437 proteins in ETBF OMVs, with 247 proteins being found in both strains. The amount of BFT-2 was similar in ETBF OMVs and NTBF+*bft* OMVs as determined by intensity-based quantification (log2 12.43 for NTBF+*bft* and log2 12.90 for ETBF). BFT-2 could not be detected by proteomic analysis in NTBF and ETBF Δ*bft* OMVs.

**Figure 3.**
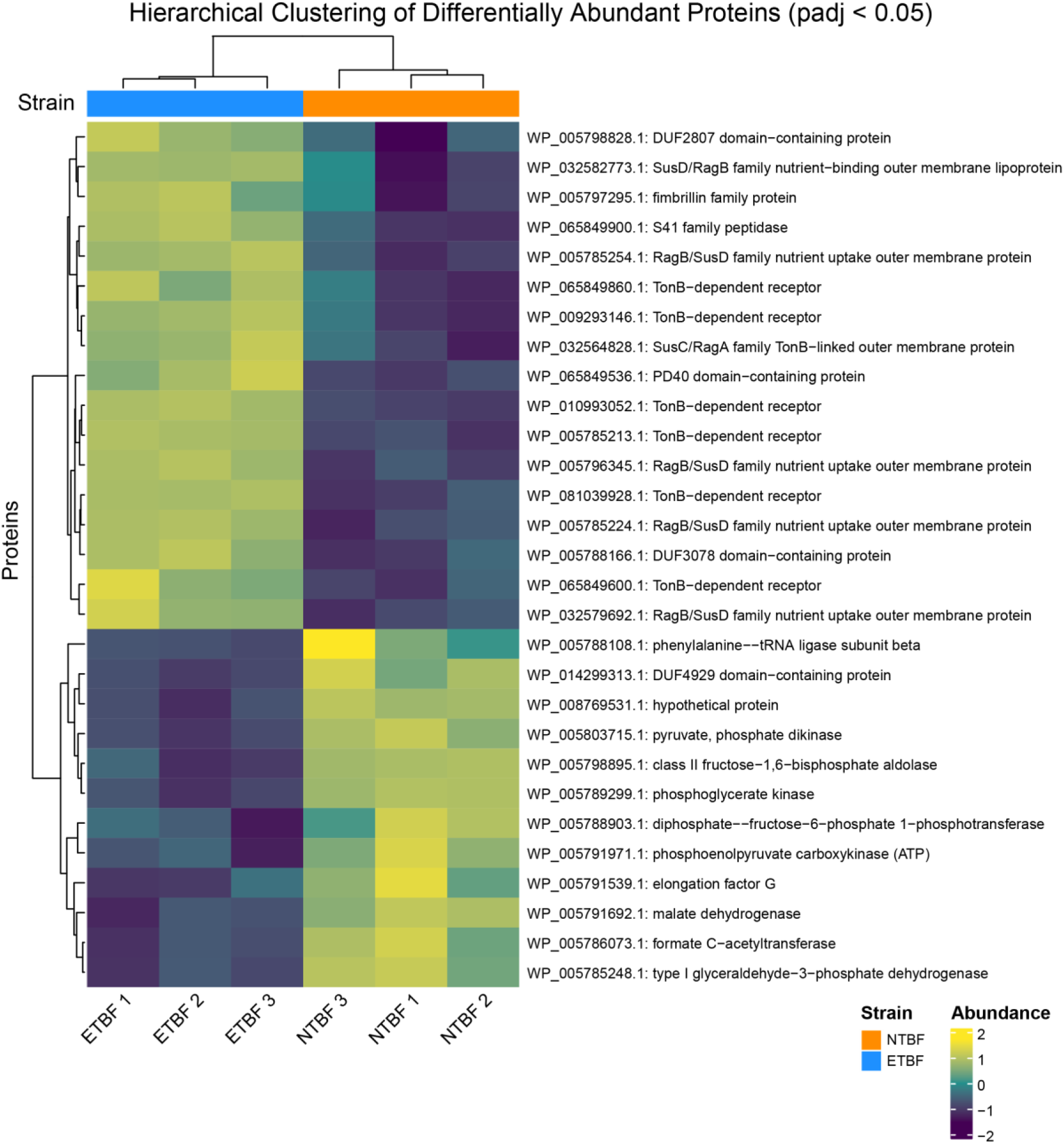
ETBF and NTBF OMV proteomes. Differential abundance analysis reveals significant differences between the proteome of ETBF and NTBF OMVs. The list of all identified proteins by proteomics analysis are presented in the Supplemental Tables. Supplemental Table 1 includes proteins detected exclusively in NTBF OMVs. Supplemental Table 2 includes proteins detected exclusively in ETBF OMVs. Supplemental Table 3 includes proteins detected in both ETBF and NTBF OMVs. Finally, Supplemental Table 4 includes differential expression of proteins in ETBF and NTBF OMVs, the results of which are depicted in this Figure. In accordance with previous research, our results show several TonB dependent receptors are more abundant in OMVs secreted by ETBF compared to NTBF strains ^36^.

### Characterization of primary human colon organoids: Development of a model to study host pathogen interactions in the colon

Biopsies of human proximal colon were obtained during clinical colonoscopy after obtaining informed consent from healthy volunteers at Dartmouth Health. As described in methods, crypt cells were dispersed and grown in Matrigel until 3D colon organoids were formed. 3D colon organoids were then seeded on Transwell filters and grown for one week to develop polarized and differentiated colon organoids. We used an imaging approach to characterize the cellular composition of colon organoids *in vitro* as monolayers. **Figures 4-5** reveal that the colon organoids contain cell types found in the proximal colon including mucin (MUC2) producing cells, CFTR expressing cells, SLC26A3 expressing cells, and enterochromaffin cells ^14,37^. These observations indicate that the human primary colon organoids are a well differentiated epithelium, have similar cell types to the colon *in vivo* and are a new model to study host pathogen interactions ^14,37^.

**Figure 4.**
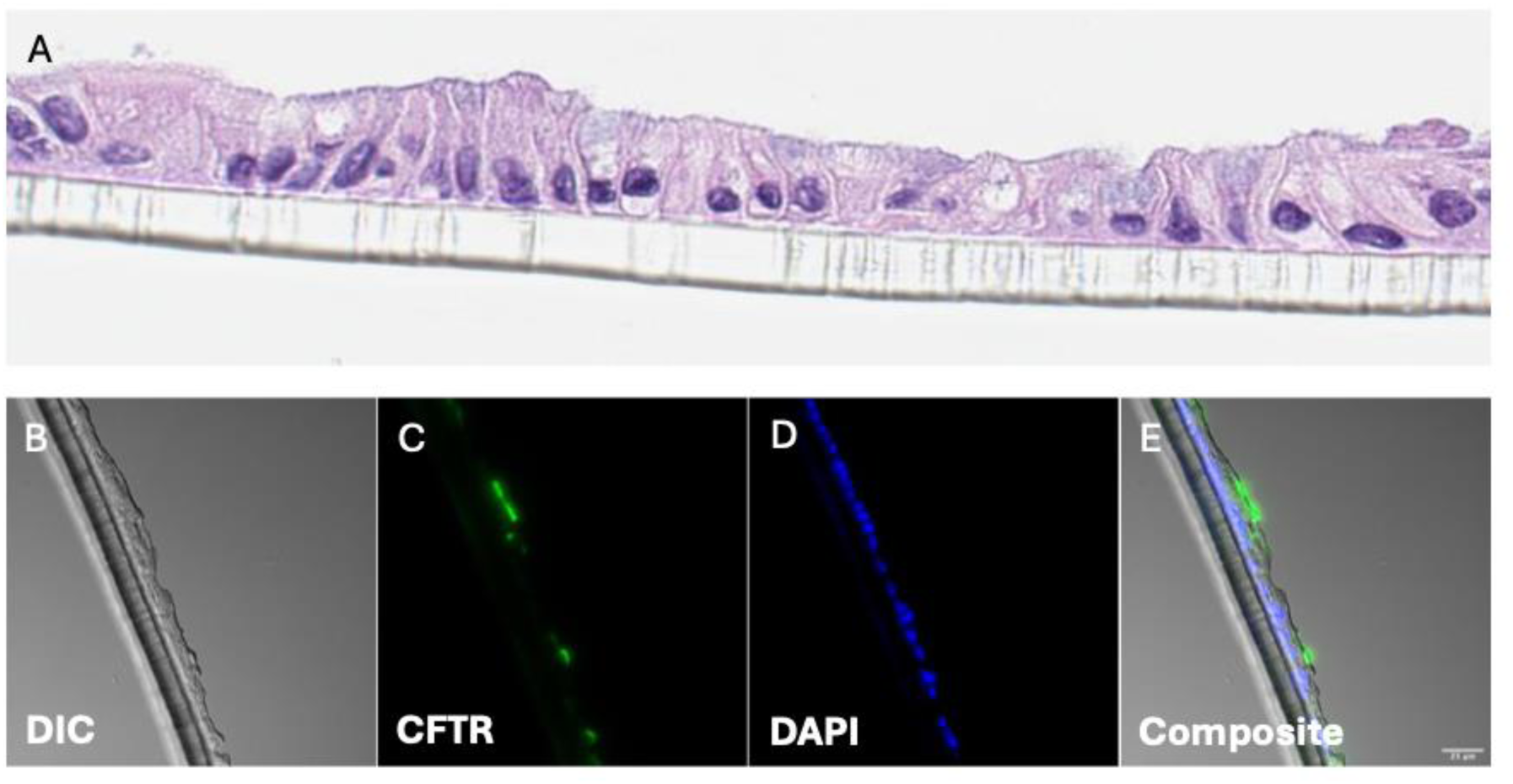
Representative images of human colon organoids. (A) Cross section of a confluent monolayer of colon organoids grown on Transwell filters (10 μm thick: H&E-stained cells). (B) differential contrast (DIC) image of a 6 μm thick frozen cross-section of differentiated colon organoids on Transwell filters. (C) Image of the same colon organoid as in B but immunofluorescent image of CFTR. (D) DAPI nuclear labeling of the same colon organoid monolayer as in B and C. (E) Composite of B-D showing apical membrane labeling of CFTR. Scale bar in panel E is 50 μm.

**Figure 5.**
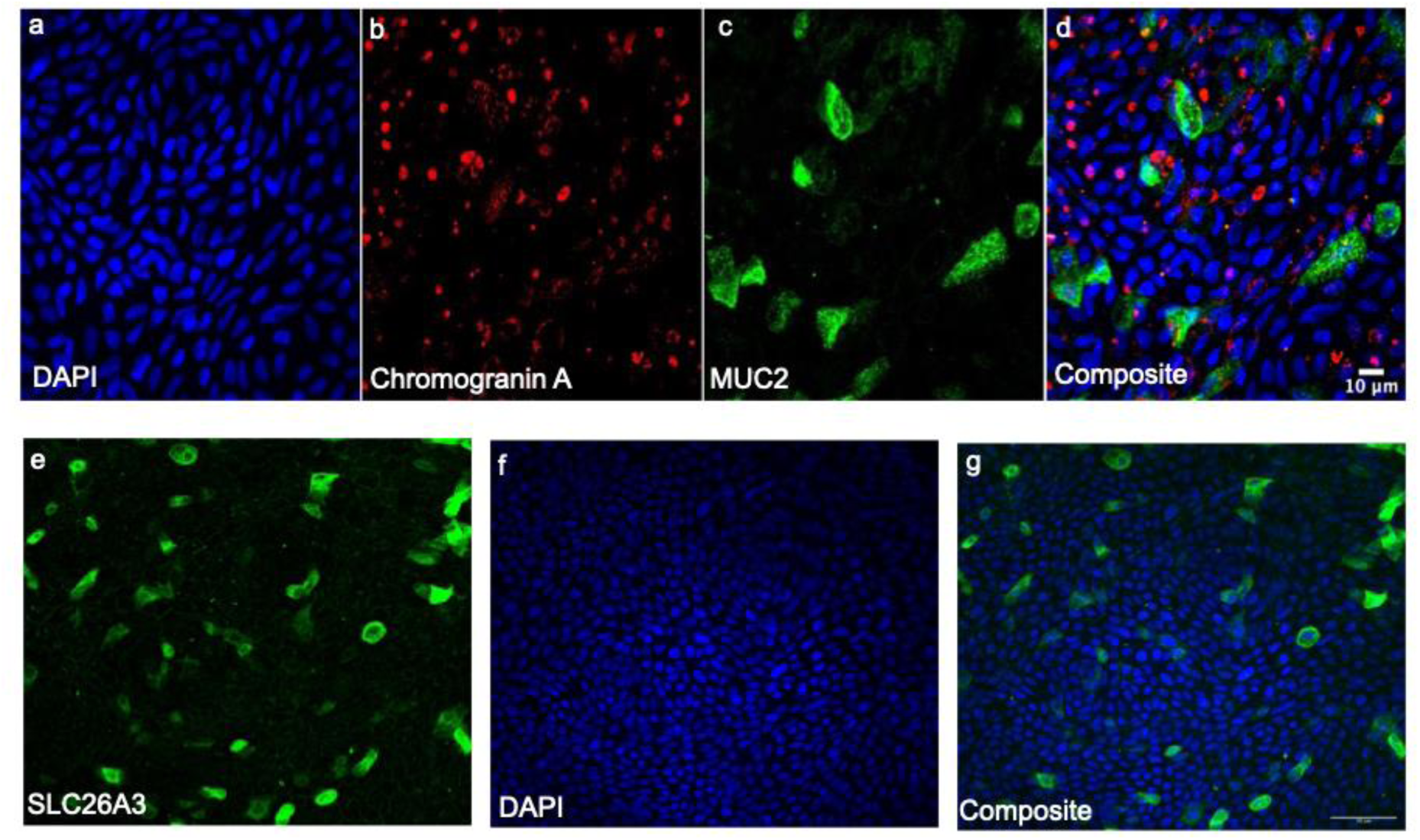
Representative *en face* images of human colon organoids on Transwell filters. (a) DAPI labeling of nuclei. (b) Chromogranin A immunofluorescence of enterochromaffin cells. (c) Immunofluorescence of MUC2. (d) Composite of A-C revealing that MUC2 and Chromogranin A are expressed in different cell types. (e) Immunofluorescence of SLC26A3 (DRA, a Cl^-^/HCO^-^ exchanger). (f) DAPI labeling of nuclei. (g) Composite of e and f. Scale bar in panel g is 50 μm.

### OMVs inhibit CFTR Cl^-^ secretion by primary human colon organoids

Previous studies have shown that recombinant BFT-2 stimulates CFTR Cl^-^ secretion, when added to the basolateral side of T84 cells ^20^. To examine the effect of OMVs secreted by *B. fragilis* on CFTR Cl^-^ secretion by primary human colon organoids, OMVs were isolated and applied to the apical side (the side of the colon exposed to Bacteroides *in vivo*) of colon organoids for 1 hr. and then forskolin-stimulated CFTR Cl^-^ secretion was measured in Ussing chambers immediately after the 1 hr. exposure, or 24 hrs. after washing the OMVs from the organoids. Forskolin increases the activity of protein kinase A, which phosphorylates and activates CFTR Cl^-^ secretion ^38^. **Figure 6** demonstrates that OMVs secreted by ETBF, ETBF Δ*bft* and NTBF+*bft* reduced forskolin stimulated CFTR Cl^-^ secretion after 1 hr. of exposure compared to process control (PC). Notably, OMVs secreted by ETBF reduced Cl^-^ secretion significantly more than OMVs secreted by NTBF that does not express BFT-2. Expression of *bft* in NTBF (NTBF+*bft*) reduced CFTR Cl^-^ secretion compared to NTBF OMVs, which do not express BFT-2.

**Figure 6.**
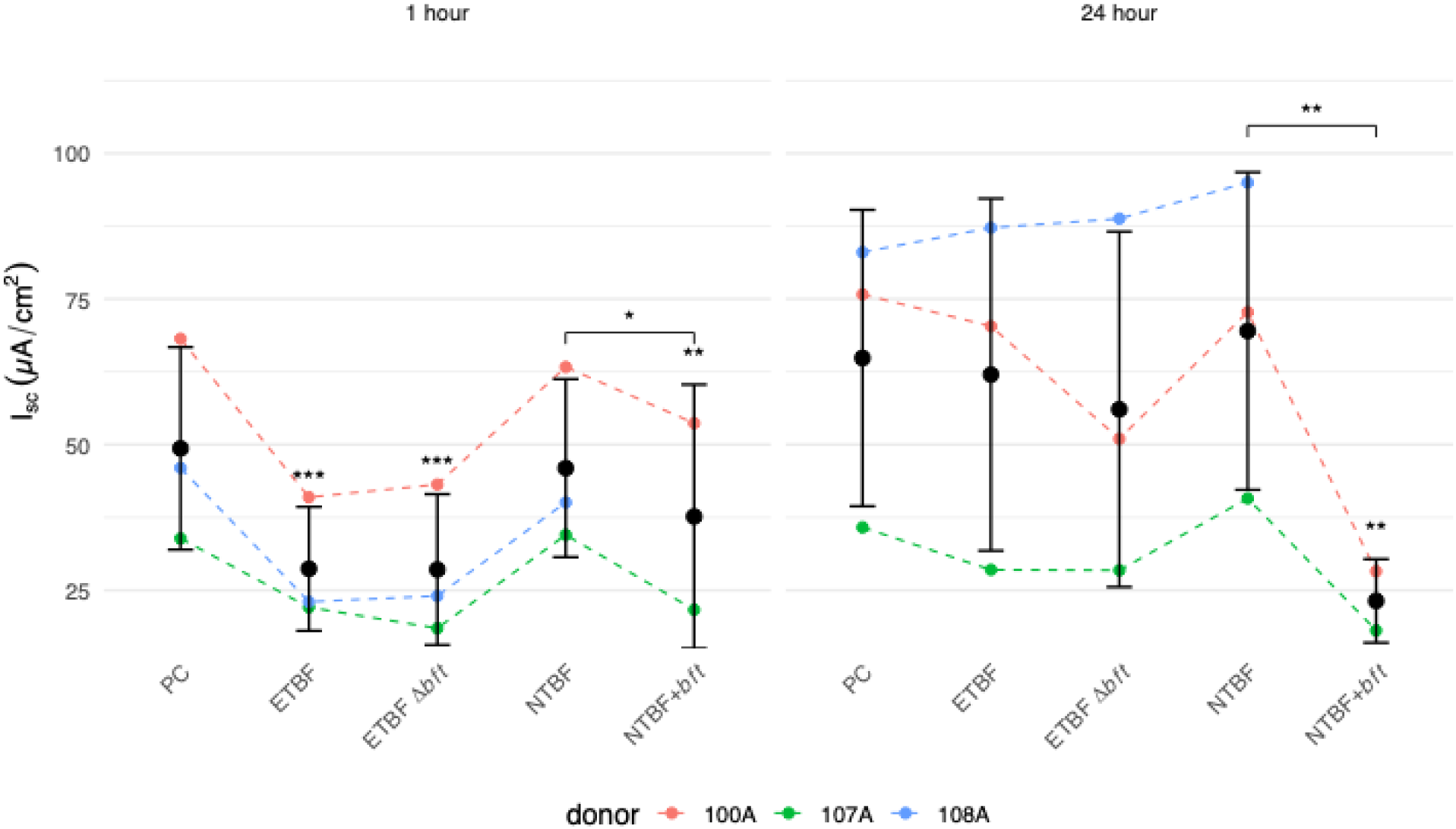
CFTR Cl^-^ secretion by human primary colon organoids measured as short circuit current. (Isc, μA/cm^2^). Left: 1 hr. after exposure to PC or OMVs (2×10^10^/ml OMVs, a concentration of OMVs similar to that measured in conditioned media and in biological fluids ^21,39^). Right: 24 hrs. after a 1 hr. exposure to PC or OMVs (2×10^10^/ml). Individual donor responses are shown as dashed colored lines, with each color representing a different donor (n = 3 donors; Donor 108A is female, Donors 100A and 107A are male). Solid black lines connect the mean current values across treatments. Data expressed as mean ± one standard deviation. Many significant differences from PC were observed at 1 hr. (ETBF, ***P = 0.00014; ETBF Δ*bft*, ***P = 0.00014; and NTBF+*bft*, **P = 0.0030). Expression of *bft* in NTBF (NTBF+*bft)* OMVs reduced CFTR Cl^-^ secretion at 1 hr. compared to NTBF OMVs, as indicated by the horizontal bar (**P = 0.01200). At 24 hrs. only NTBF+*bft* OMVs reduced CFTR Cl^-^ secretion compared to PC (P=0.008). The horizontal bar indictes that NTBF+*bft* OMVs significantly reduced CFTR Cl^-^secretion (**P = 0.004) compared to NTBF OMVs. Statistical analysis was performed using mixed-effects linear models with donor as a random effect for each time point separately. Experiments done with three donors of colon cells, three technical replicates per donor and three different preparations of OMVs. Technical replicates were averaged as a single data point.

Twenty-four hrs. after a 1 hr. exposure to OMVs, CFTR Cl^-^ secretion was not significantly reduced by ETBF, ETBF Δ*bft*, or NTBF when compared to PC (**Figure 6**). However, 24 hrs. after exposure to NTBF+*bft* OMVs, CFTR Cl^-^ secretion was less that CFTR Cl^-^ secretion compared to NTBF OMVs (**Figure 6**).

OMVs had no effect on the amiloride-sensitive short circuit Na^+^ currents (Isc, μA/cm^2^), either after a 1 hr. exposure to OMVs or after 24 hrs. following the 1 hr. exposure to OMVs (**Figure 7**).

**Figure 7.**
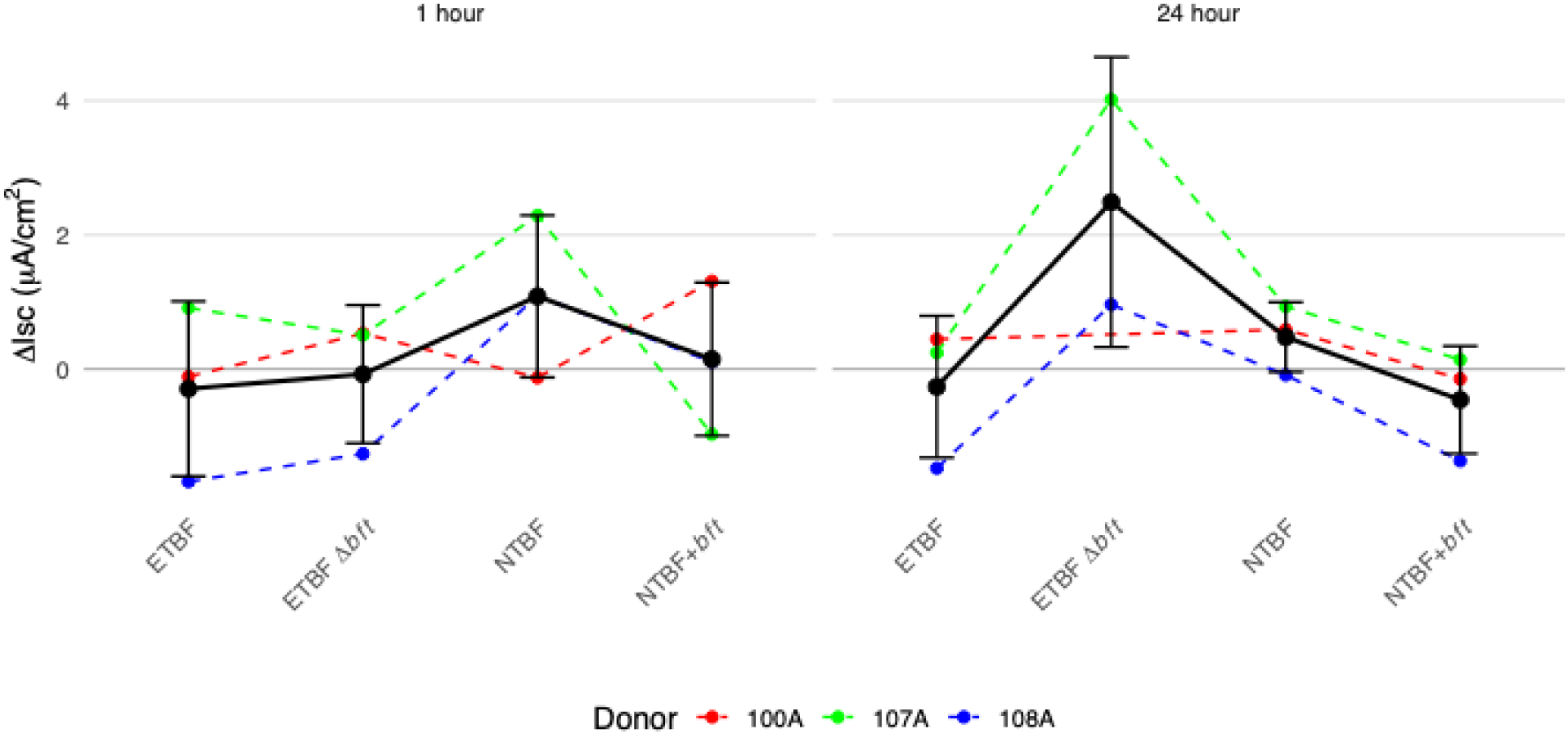
OMVs did not modify amiloride-sensitive short circuit Na^+^ currents. Amiloride-sensitive Na^+^ currents in response to OMVs (2×10^10^/ml OMVs) compared to PC. The difference in amiloride-sensitive currents (ΔIsc) between PC and OMVs exposed colon organoids is plotted. Colon organoids were exposed to OMVs for 1 hr. or 24 hrs. Each measurement of amiloride-sensitive current for each donor was adjusted by subtracting the Process Control (PC) current from the same Ussing chamber session and the same donor. Data from each donor (100A = red, 107A = green, 108A = blue) is depicted with dashed lines. The Isc OMVs vs. PC) for all donors is shown with a solid black line with ± one standard deviation. The horizontal line at zero represents no change from PC. Statistical analysis was performed using mixed-effects linear models with donor as a random effect for each time point separately. Experiments done with three donors of colon cells, three technical replicates per donor and three different preparations of OMVs. Technical replicates were averaged as a single data point.

Experiments were also conducted to determine if OMVs had an effect on TER, since previous studies have shown that recombinant BFT-2 reduced TER in several colon cell lines ^9,15,17,19,20^. However, none of the OMVs had a significant effect on TER (**Figure 8**) compared to process control (PC) at 1 hr. and 24 hrs.

**Figure 8.**
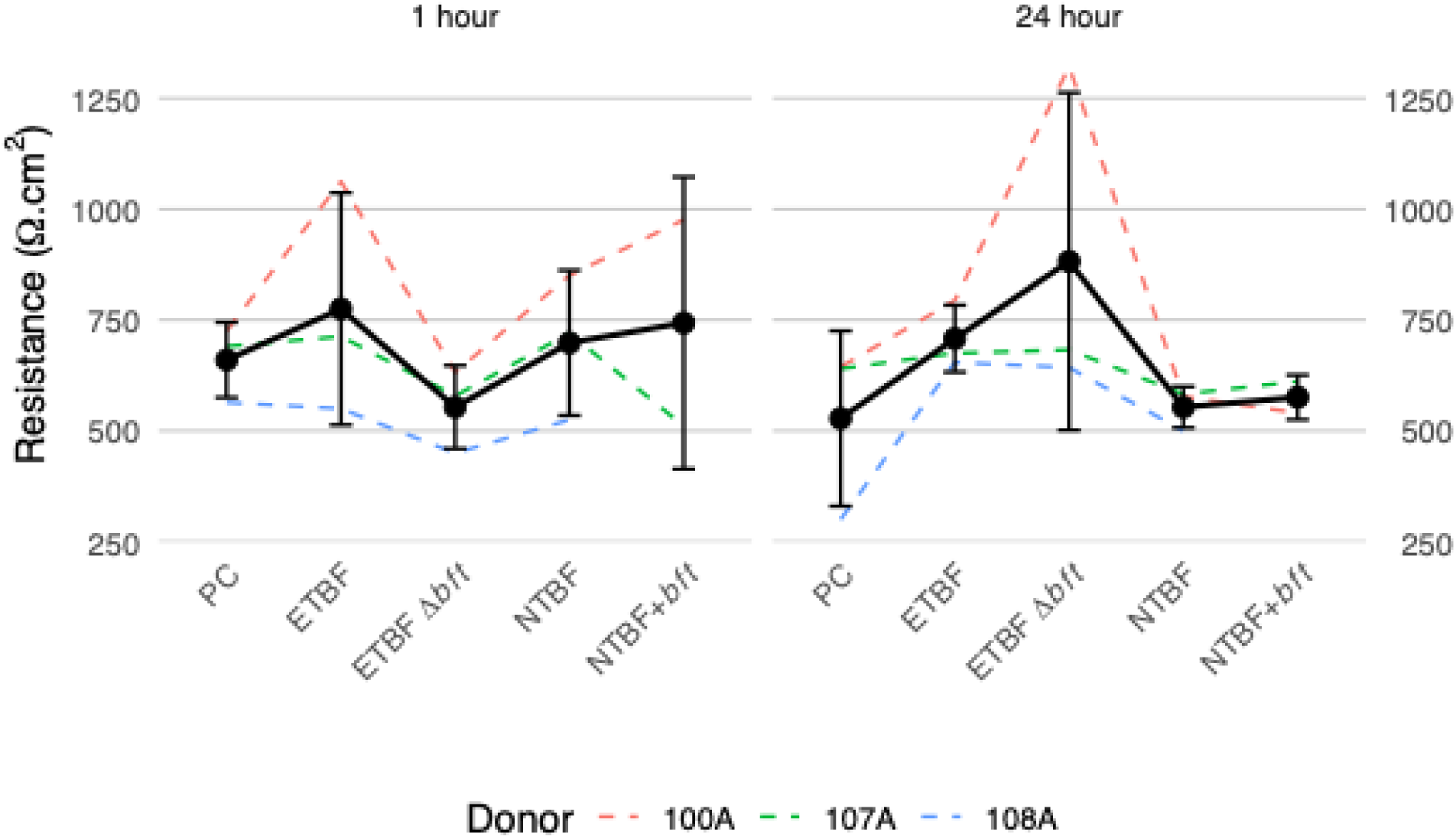
Transepithelial resistance (TER) of primary human colon organoids. TER was measured in colon organoids in which CFTR Cl^-^ secretion was measured 1 hr. (left figure) or 24 hrs. (right figure) after exposure to PC or OMVs (2×10^10^/ml OMVs) secreted by ETBF, ETBF Δ*bft*, NTBF and NTBF+*bft*. Individual donor responses are shown as dashed colored lines, with each color representing a different donor. Solid black lines connect the mean TER values across treatments. Data expressed as mean ± one standard deviation. Statistical analysis was performed using mixed-effects linear models with donor as a random effect for each time point separately. Experiments done with three donors of colon cells, three technical replicates per donor and three different preparations of OMVs. Technical replicates were averaged as a single data point. No significant differences compared to PC (1hr. and 24 hr. exposure) were observed (P>0.05).

### Bacteroides reduces CFTR Cl^-^ secretion by T84 cells but has no effect on TER

We also conducted studies on T84 cells, a colon cell line that has been used to study the effects of recombinant BFT-2 ^40–42^. Previous studies have shown that recombinant BFT-2 reduces the TER in several colon cell lines ^9,15,17,19,20^. To determine if *B. fragilis* OMVs also reduced TER and influenced CFTR Cl^-^ secretion, we conducted Ussing experiments on monolayers of T84 cells under conditions identical to the previously described exposure of primary human colon organoids. Compared to PC, CFTR Cl^-^ secretion was reduced by all OMVs (**Figure 9a**). ETBF OMVs were not more effective than NTBF OMVs in reducing CFTR Cl^-^ secretion. Moreover, deletion of *bft-2* in ETBF did not significantly reduce the effect of ETBF OMVs on CFTR Cl^-^secretion. OMVs also had no effect on TER (**Figure 9b**). Thus, OMVs reduced CFTR Cl^-^secretion by both primary human colon organoids and T84 cells, without altering TER.

**Figure 9.**
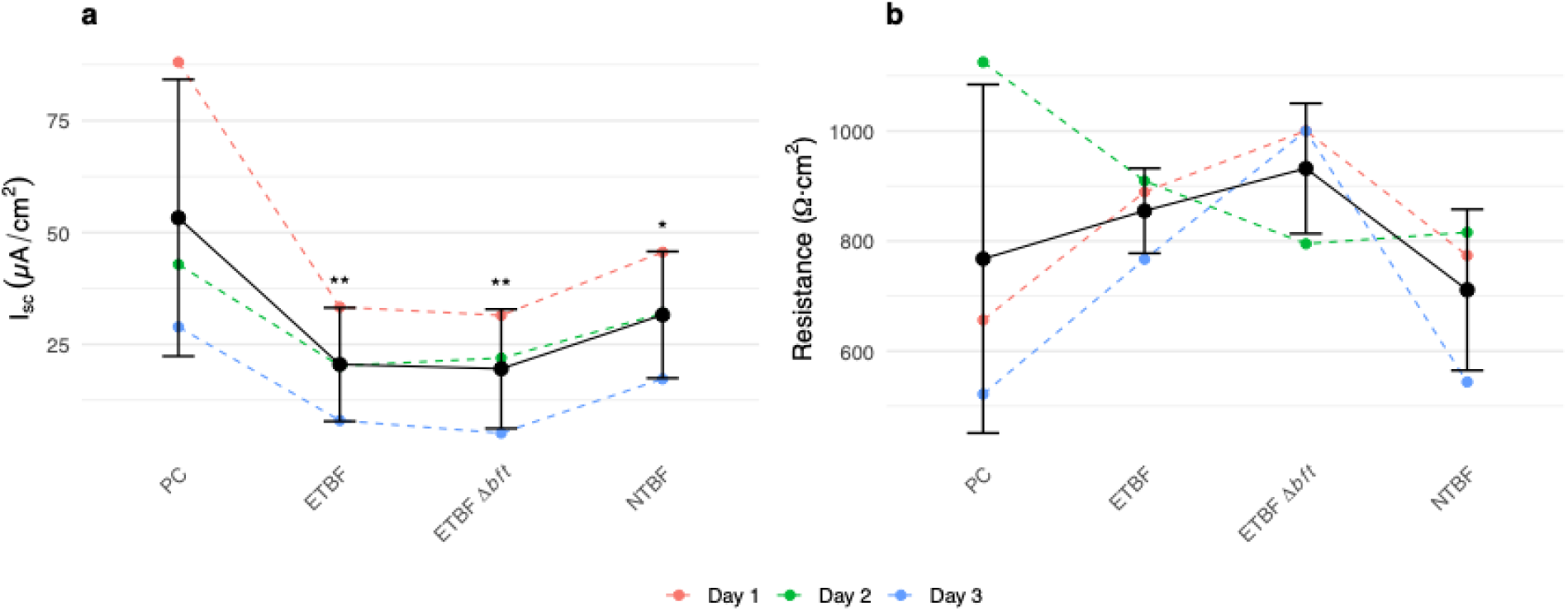
CFTR Cl^-^ secretion and TER by T84 cells exposed to OMVs. The protocol was identical to that performed on primary human colon organoids. T84 monolayers were exposed to OMVs (2×10^10^/ml) for 1 hr. and CFTR Cl^-^ secretion (short-circuit current: Isc) was measured. Data represent individual experiments (dashed lines) grouped by experimental date (colors) with means ± one standard deviation. (a) CFTR Cl^-^ secretion. Each line connects experiments conducted on the same day on the same pass of T84 cells. Compared to PC, CFTR Cl⁻ secretion was significantly reduced by ETBF (**P = 0.006), ETBF Δ*bft* (**P = 0.005), and NTBF (*P = 0.032) OMVs. (b) TER in the same set of T84 cells used for measurements of CFTR Cl^-^ secretion. No significant differences in TER compared to PC were observed after 1 hr. of exposure to OMVs. Statistical analysis was performed using mixed-effects linear models with donor as a random effect for each time point separately. Experiments done with three different preparations of OMVs. Technical replicates were averaged as a single data point.

### OMVs do not alter tight junction or cell adhesion protein abundance

Previous studies have also shown that recombinant BFT-2 rapidly (<1 hr.) reduced the expression of the cell adhesion protein E-cadherin in cell lines ^9,15,17,19,20^. However, we observed no effect of Bacteroides OMVs on the abundance of E-cadherin or on the tight junction protein claudin-2 in human colon organoids as determined by immunofluorescence microscopy (**Figure 10**).

**Figure 10.**
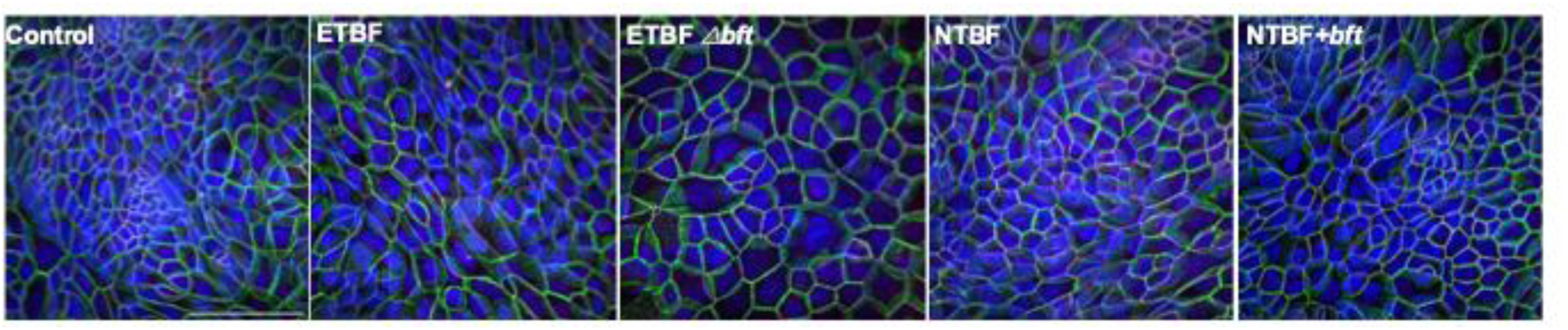
Representative composite images of human colon organoids. Organoids were grown on Transwell filters and exposed to OMVs (2×10^10^/ml) for 1 hr. Claudin-2 (magenta) and E-cadherin (green) were identified by immunofluorescence. DAPI (blue) identified nuclei. Each image is a composite of colon organoids labeled for claudin-2, E-cadherin and nuclei (DAPI). Confocal images were analyzed by an investigator who was blinded to the treatment. There was no difference in claudin-2 or E-cadherin abundance using confocal microscopy and the Fiji open-source image processing package to quantify fluorescence intensity (see Figure 11 for data).

Studies were also conducted to measure E-cadherin and claudin-2 fluorescence imaged by confocal microscopy. None of the OMVs had a significant effect on E-cadherin or claudin-2 fluorescence (**Figure 11**) confirming the immunofluorescent images in Figure 10.

**Figure 11.**
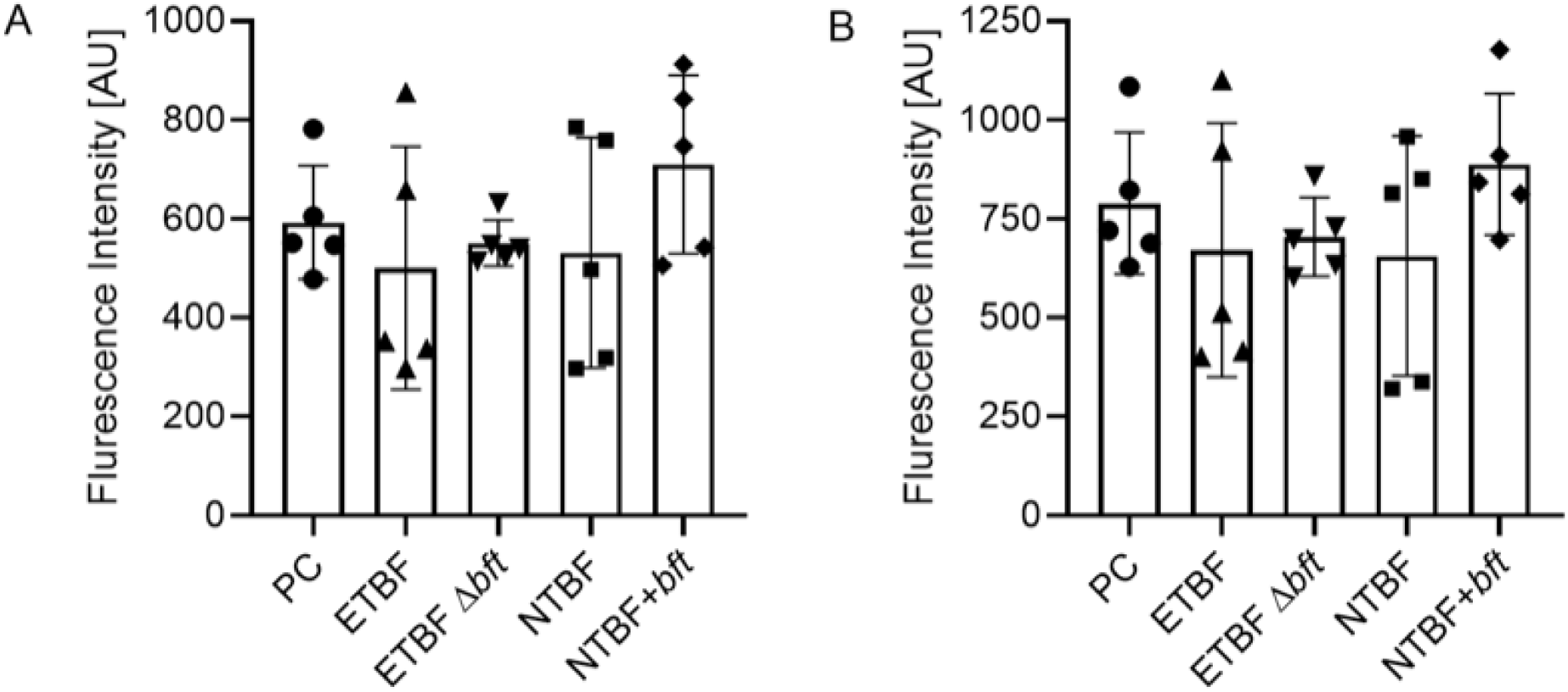
Measurements of E-cadherin (A) and claudin-2 (B) epifluorescence. Data were generated by a co-author blinded to the experimental treatment. Five random images were chosen for each treatment. There was no significant effect of OMVs (2×10^10^/ml) on E-cadherin or claudin-2 abundance. AU, arbitrary fluorescent units. Data expressed as mean ± one standard deviation.

### Cytokine secretion by human primary colon organoids

Previous studies on cell lines and human colon tissue reported that *B. fragilis* and recombinant BFT-2 stimulates IL-8 secretion ^9,15,17–20^. To examine the effect of OMVs on the innate immune response, colon organoids were grown on Snapwell filters and exposed to PC or OMVs in the apical media for 1 hr. or 24 hrs. after OMVs were removed by washing. **Figure 12** demonstrates that OMVs had no significant effect on IL-8 secretion compared to PC. Moreover, there was no significant effect of any OMVs compared to PC on the secretion of 48 cytokines measured using the MILLIPLEX MAP Human Cytokine/Chemokine 48-Plex cytokine assay, although there were some trends in the response. For example, compared to PC, 24 hrs. after exposure to ETBF OMVs, and NTBF+*bf*t OMVs IL-8 increased compard to PC; however the changes were not statistically significant.

**Figure 12.**
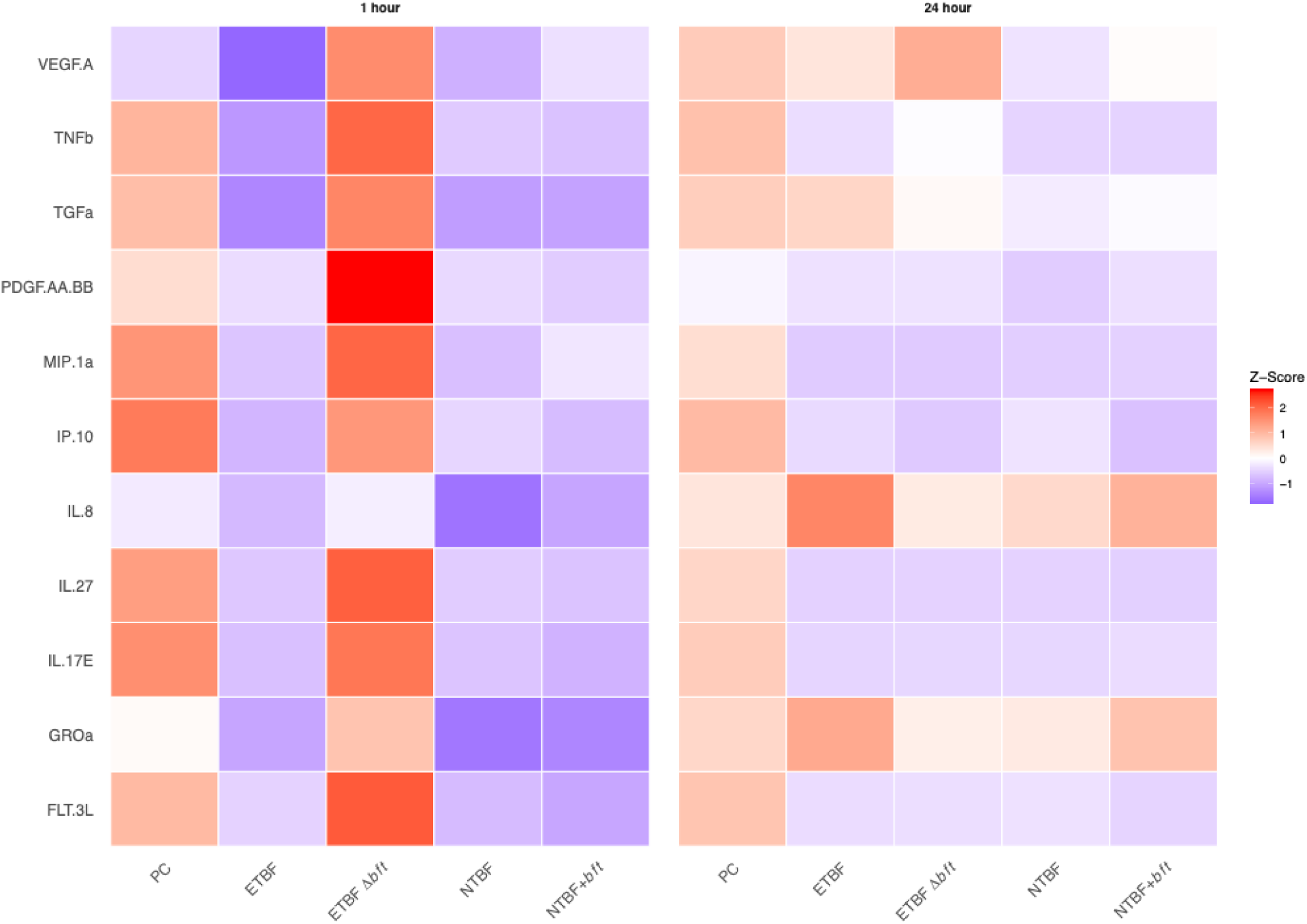
Cytokine secretion following 1 hr. and 24 hrs. after exposure to OMVs secreted by *Bacteroides fragilis* or to PC. There were no significant effects of OMVs (2×10^10^/ml) versus PC. Values represent means across all donors (n = 3 donors) for each treatment condition.

Color scale indicates standard deviations from the time point mean: red represents above-average expression, blue represents below-average expression, and white represents average expression for each cytokine at each time point. Only cytokines with complete data (values above detection) across all conditions are shown. Data were log2-transformed prior to z-score calculation.

## Discussion

There were two primary goals of this study: (1) To develop a well-differentiated primary culture of human colon organoids and (2) to elucidate the effect of endogenous BFT-2 in OMVs secreted by *B. fragilis* on human primary colon organoids. We sucessfully developed a model of colon organoids from three donors that displayed a differentiated epithelium containing cell types found in the proximal colon including mucin (MUC2) producing cells, CFTR expressing cells, SLC26A3 (DRA, a Cl^-^/HCO^-^ exchanger that plays a key role in fluid absorption and enterocyte acid/base balance) expressing cells, and enterochromaffin cells that underlie epithelial cells and secrete serotonin, which stimulates GI motility and fluid and electrolyte secretion ^14,37^. These observations indicate that the human primary colon organoids are a representative model to study host pathogen interactions in the human colon ^14,37^. Studies were also conducted to examine the effect of OMVs on CFTR Cl^-^ secretion, amiloride-sensitive sodium absorption, TER, and cytokine secretion. We found that OMVs secreted by most strains of Bacteroides significantly reduced CFTR Cl^-^ secretion by colon organoids compared to PC after 1 hr. exposure. Deletion of the *bft-*2 gene from ETBF reduced the inhibitory effect of ETBF OMVs on CFTR Cl^-^ secretion. Moreover, NTBF+*bft* OMVs were significantly more inhibitory to CFTR Cl^-^ secretion compared to OMVs secreted by NTBF. Both observations indicate that BFT-2 inhibits CFTR Cl^-^ section. Twenty-four hrs. after exposure to OMVs, only NTBF+*bft* OMVs reduced CFTR Cl^-^ secretion compared to NTBF OMVs and PC. Thus, BFT-2 in OMVs was, at least in part, responsible for the decrease in CFTR Cl^-^ secretion by primary human colon organoids. Since OMVs secreted by strains of *B. fragilis* that lacked *bft*-2 (BFT Δ*bft* and NTBF) also reduced CFTR Cl^-^ secretion, there must be other factors in OMVs including, but not limited to, proteins or miRNAs that inhibit forskolin-stimulated CFTR Cl^-^ secretion. By contrast to results from human colon organoids all OMVs secreted by *B. fragilis* inhibited CFTR Cl^-^ secretion by T84 cells to the same extent. The reason for this difference between the effect of OMVs on T84 cells and human colon organoids is unknown, but it may be related to several factors, including, but not limited to, possible differences in the time dependent effects of BFT-2 as well as differential expression of gene and signaling pathways between T84 tumor cells and primary human colon organoids. Additional studies are required to identify these factors.

The observation that OMVs inhibited CFTR Cl^-^ secretion by primary human organoids is interesting to note since infection with ETBF causes secretory diarrhea, which one might expect to be caused by increased CFTR Cl^-^ secretion, a primary driver of fluid secretion by the colon^13,14^. However, there is differential expression and regulation of ion transporter proteins along the length of the colon, as well as differential responses to drugs, hormones, and immune factors along the length of the colon ^13,14^. Accordingly, our results are applicable to the proximal human colon, and other segments of the colon may react differently to OMVs and BFT-2. It is the composite effect of ETBF and BFT-2 on the gastrointestinal tract that determines the response *in vivo* to ETBF.

Our studies also did not observe a significant effect of OMVs on cytokine secretion by human colon organoids. Most notably, IL-8 secretion was not significantly increased by OMVs(**Figure 12**), in contrast to studies using recombinant BFT-2 in cells lines reporting that recombinant BFT-2 increased IL-8 secretion ^9,15,17–20^. The reason for this difference between the effect of OMVs on colon cell lines and human colon organoids is unknown, but it may be related to several factors, including, but not limited to, differences in the amount of recombinant versus native BFT-2 in OMVs, as well as differences in gene signaling pathways between colon cell lines and human primary colon organoids. Studies have also shown that the effect of recombinant BFT-2 and OMVs on colon cell lines is dose dependent, and it is likely that the concentration of BFT-2 in OMVs is less that the doses of recombinant BFT-2 utilized in studies on cell lines ^11,18^. We conclude that biologically relevant levels of OMVs^21,39^ secreted by *B. fragilis* containing BFT-2 inhibit CFTR Cl^-^ secretion, but have no effect on amiloride-sensitive sodium reabsorption, claudin-2, E-cadherin or TER, on primary human colonoids.

Fragipain was not detected in any of the OMV samples (Supplemental Table 4). Fragipain is known to cleave BFT-2 into its active form ^43^, however, studies suggest that this protein is not required for cleavage of BFT-2 in antibiotic treated mice ^44^. More specifically, fragipain is unnecessary when recombinant BFT-2 is mixed with the colon mucus of pathogen free or germ-free mice, suggesting the mucosal activation process may be more prevalent in gastrointestinal conditions containing increased mucus, such as CF ^44^. Notably, primary human colon organoids also secrete mucin. PBS is also able to facilitate the cleavage of BFT-2 into its active form, and further suggests that fragipain is not required for BFT-2 activation ^44^.

In accordance with previous research, our proteomics data revealed that several TonB dependent receptors are more abundant in OMVs secreted by ETBF compared to NTBF strains (Supplemental Table 4) ^45^. TonB dependent receptors are one part of outer membrane structures that are critical for the influx of nutrients via SusC and SusD outer membrane proteins, which were elevated in ETBF OMVs compared to NTBF OMVs ^36^. An increase in abundance of these proteins suggests an enhanced need for ETBF to increase nutrient acquisition, thereby improving its ability to compete for similar niches that NTBF does not struggle to fill. Regardless, these results pose interesting questions for mechanistic follow up experiments in the future.

Importantly, although several studies have shown that recombinant BFT-2 degrades E cadherin and claudin-2 in cell lines^9,15,17,19,20^, none of the OMVs tested herein had any effect on the abundance of the tight junction protein claudin-2, the cell adhesion protein E-cadherin, or on the TER. Claudin-2 is an important component of the tight junction, and reductions in claudin-2 abundance decrease TER^46,47^. Several possible explanations may account for the difference in the TER, E-cadherin, claudin-2, and cytokine response of colon tumor cell lines and primary human colon organoids. First, as noted above, the effects of BFT-2 are dose dependent ^18^.

OMVs secreted by ETBF and NTBF+*bft* contain endogenous levels of BFT-2. It is possible that the concentration of recombinant BFT-2 in previous studies was much higher than the amount of native BFT-2 in OMVs, although a comparison of concentrations is not possible given available data. Second, the duration of exposure to OMVs may be a factor. The literature reports that the effects of recombinant BFT-2 on E-cadherin and claudin-2 in cell lines required anywhere from 1 hr. to several hrs. of exposure^3,9,10^. Our study was limited to a one-hr. exposure, which has been long enough to produce significant effects in many previous studies with recombinant BFT-2 and colon cell lines ^3,9,10^.

There are a few other reports describing the development of monolayers of epithelial cells derived from healthy colon organoids ^31,32^. Sen et al. ^32^ reported that the gene expression profile of human colon organoids was significantly different from the Caco-2 cell line, an immortalized cell line of human colorectal adenocarcinoma cells, which had a gene expression profile similar to the small intestine. They reported that the effect of *Bifidobacterium longum* on gene expression was not conserved between Caco-2 and colon organoids. Taglieri et al ^31^ reviewed the literature on human colon organoids and noted that air liquid interface culture increased differentiation of colon organoids. However, none of the other studies on colon organoids exposed them to OMVs or reported CFTR Cl^-^ secretion.

OMVs secreted by most strains of *B. fragilis* in the present study inhibited forskolin-stimulated CFTR Cl^-^ secretion by the proximal colon and T84 cells, which would reduce salt and fluid secretion by the colon. However, studies in animals report that recombinant BFT-2 induces secretory diarrhea ^3,6–8^. One possible reason for the different effects of recombinant BFT-2 and OMVs containing BFT-2 is that the effect of BFT-2 is dose dependent, and that recombinant BFT-2 levels in published studies may be higher than the abundance of BFT-2 in OMVs. As demonstrated in Figure 3 and the Supplemental Tables, OMVs contain many proteins in addition to BFT-2, which may influence CFTR Cl^-^ secretion by primary human colon organoids. Finally, it is important to note that there are other pathways that mediate transepithelial salt and water transport by the proximal colon, including electroneutral NaCl and fluid reabsorption mediated by the parallel operation of Na^+^/H^+^ and Cl^-^/HCO_3_ ^-^ (SLC26A3) exchangers. Since Ussing chamber studies only measure electrogenic transport, it is possible that recombinant BFT-2 (and endogenous BFT-2) may inhibit SLC26A3 and/or the Na^+^/H^+^ exchanger *in vivo* and thereby reduce electroneutral NaCl and fluid reabsorption by the colon. Such an effect may contribute to the ETBF-induced secretory diarrhea, despite the OMV reduction in electrogenic CFTR Cl^-^ secretion reported herein. Thus, extensive additional studies, beyond the goals of the present study, are required to determine if BFT-2 and OMVs inhibit Na^+^/H^+^ and Cl^-^/HCO_3_ ^-^ electrically neutral NaCl and fluid reabsorption in primary human colon organoids. Such studies would require use of radioactive isotopes and chemical inhibitors of electrically neutral transporters and/or antisense experiments to selectively reduce electroneutral transporter expression.

We also did not observe an effect of any OMVs on the TER or epithelial tight junction or cell adhesion structure. In many, but not all, previous studies in colon cell lines, recombinant BFT-2 quickly (< 1 hr.) reduced the abundance of E-cadherin and the TER ^9,15,17,19,20^. Although the reason for this difference is unknown, one possible explanation for the different effects of recombinant BFT-2 and OMVs is that, as noted above, the effect of recombinant BFT-2 is dose and time dependent, and that recombinant BFT-2 levels in published studies may be higher than the abundance of endogenous BFT-2 in in OMVs.

In contrast to studies using cells lines and recombinant BFT-2 ^10,48–50^, we did not observe a significant increase in IL-8 secretion in primary human colon organoids exposed to OMVs. The lack of effect of BFT-2 and OMVs on IL-8 secretion in our study are consistent with the observation that OMVs did not reduce E-cadherin, an effect reported to be required for BFT-2 to stimulate IL-8 secretion ^10,48–50^. OMVs did not affect the secretion of any of the 48 cytokines as determined by the MILLIPLEX MAP Human Cytokine/Chemokine 48-Plex cytokine assay. The reason for this difference is also unknown and requires additional studies.

There are a few limitations of our study. First, since there are many differences in the content of OMVs secreted by ETBF and NTBF such as DNA, RNA, miRNA, proteins (this study) and other virulence factors ^24,51^, in addition to BFT-2, it is possible that some of these factors may act alone of in concert with BFT-2 to inhibit CFTR Cl^-^ secretion. Second, although we did not complement the *bft-2* gene in ETBF Δ*bft*, due in part to the technical challenge of complementation in our clinical strain, we have complementary lines of evidence to support our conclusions that BFT-2 inhibits CFTR Cl^-^ secretion in colon organoids. First, we utilized two strains o*f B. fragilis* that lack BFT-2 (ETBF Δ*bft* and NTBF) and two that secrete BFT-2 (ETBF and NTBF+*bft*). Our results on primary human colon organoids are similar in ETBF Δ*bft* and NTBF, as are the results using ETBF and NTBF+*bft*.

Our study has numerous strengths. First, our experiments were conducted on well differented human colon organoids from three healthy donors, and a cell line (T84). OMVs inhibited CFTR Cl^-^ secretion by colon organoids and T84 cells, but had no effect on TER, or the expression of claudin-2 and E-cadherin. Second since *B. fragilis* resides in mucus overlying colon organoids, OMVs containing BFT-2 are biologically relevant. Notably, the concentration of OMVs used in the present study was similar to levels observed in conditioned cell culture media and biological fluids^21,39^, and the same number of OMVs was used in all experimental groups ^21,39^.

In conclusion, we have examined the effects of OMVs on well differentiated primary cultures of human colon organoids secreted by *B. fragilis* containing *bft-2 (*ETBF and NTBF+*bft)*, a strain of *B. fragilis* in which *bft-2* was knocked out (ETBF Δ*bft*), and an NTBF strain that does not express *bft-2.* In all cases, BFT-2 containing OMVs secreted by *B. fragilis* reduced forskolin-stimulated CFTR Cl^-^ secretion by colon organoids more than the same strain lacking *bft-2* but had no effect on tight junction structure (E-cadherin and claudin-2), TER, amiloride-sensitive sodium absorption, or IL-8 secretion. Similar experiments on the T84 cell line also revealed that OMVs secreted by *B. fragilis* reduced CFTR Cl^-^ secretion and had no effect on TER. We conclude that *B. fragilis* OMVs containing biologically significant levels of BFT-2 inhibit forskolin stimulated CFTR Cl^-^ secretion by primary human colon organoids, without effecting TER, amiloride-sensitive Na^+^ currents and cytokine secretion. Additional studies, beyond the scope of the present study, are required to elucidate the cellular mechanisms whereby OMVs rapidly inhibit CFTR Cl^-^ secretion. Although the results in this study contradict many studies using cell lines and recombinant BFT-2, we anticipate that our results will stimulate a discussion of the effects on BFT-2 on the colon.

## METHODS

### Bacteroides culture and genetics

*B. fragilis* NCTC 9343 was chosen as the non-toxigenic *B. fragilis* (NTBF) strain, since it is used widely ^52–54^. *B. fragilis* 86-5443-2-2 was chosen as the enterotoxigenic *B. fragilis* (ETBF) strain for similar reasons, and because it expresses BFT-2, which is the most virulent of the three BFT isotypes ^52–54^. *B. fragilis* 86-5443-2-2 with a *bft* deletion (ETBF Δ*bft*), provided by Dr. Cynthis Sears, was confirmed by PCR ^55^. To construct the NTBF+*bft* strain, we first PCR amplified *bft-2* from *B. fragilis* 86-5443-2-2 and cloned it via Gibson Assembly into the SalI and NcoI sites of the *Bacteroides* expression plasmid pNBU2-P1T-DP-A21 ^56^. This plasmid integrates in single copy into the chromosomal att sites of *Bacteroides fragilis.* The sequence of the resulting plasmid was confirmed by Sanger sequencing before transformation into *E. coli* S17-1 cells and conjugation via overnight aerobic solid agar co-culture mating into *B. fragilis* 9343 (NTBF). Following incubation, matings were resuspended and plated on BHIS agar containing gentamycin and erythromycin. Att chromosomal integration site was confirmed by PCR ^57^. Bacteroides strains were grown on supplemented brain heart infusion medium (BHIS) agar plates or liquid broth cultures supplemented with hemin and 1μg/ml of vitamin K3 (Acros Organics/Fisher Scientific, Geel, Belgium) under anaerobic conditions (10% CO2, 10% H2, 80% N2) in a Whitley A55 anaerobic chamber (Don Whitley Scientific, Victoria Works, UK) at 37°C. All BHIS agar and liquid media contained 60 μg/ml of gentamycin sulfate. For OMV purification, *B. fragilis* strains were streaked on BHIS agar plates and were grown in anaerobic conditions for 48 hrs. Single colonies were used to inoculate 5mL of BHIS liquid medium cultures, which were grown anaerobically for 24 hrs. The cultures were then expanded into 200mL BHIS cultures using a 1:100 dilution. The final cultures were grown for 24 hrs. prior to being utilized to isolate OMVs.

### Outer membrane vesicle (OMV) isolation

OMVs were isolated as described by us and others previously ^23,58^. Briefly, all overnight culture supernatants were centrifuged for 1 hr. at 2800 g and 4°C to pellet bacteria. The supernatant was filtered twice through 0.45 μm PVDF membrane filters (Millipore, Billerica, MA, USA) to remove any contaminating bacteria and concentrated with 30K Amicon filters (Millipore, Billerica, MA, USA) at 2800 g and 4°C to obtain ∼200μL of concentrate. The concentrate was resuspended in OMV buffer (20 mM HEPES, 500 mM NaCl, pH 7.4) and subjected to ultracentrifugation using a TH-660 Swinging Bucket rotor (Thermo Fisher) for 2 hrs. at 200,000 g and 4°C to pellet OMVs. OMV pellets were then re-suspended in 60% OptiPrep Density Gradient Medium (Sigma-Aldrich, Cat. # D1556) and layered with 35%, 30% and 20% OptiPrep diluted in OMV buffer. OMVs in OptiPrep were centrifuged for 18 hrs. at 100,000 g and 4°C. 500μl fractions were taken from the top of the gradient, with OMVs residing in fractions 5-8, as determined by proteomic analysis of OMVs.

### Electron microscopy of OMVs

OMVs were visualized with negative staining transmission electron microscopy as described previously ^22^. The microscopist was blinded to the experimental treatment to minimize observer bias. Briefly, an aliquot of each sample was applied onto a freshly glow-discharged 200-mesh nickel grid and allowed to sit for two minutes. After incubation, excess fluid was gently removed with filter paper, and the grids were sequentially rinsed with Millipore-filtered distilled water. While the grids remained slightly damp, they were placed onto a drop of Nano-W tungsten stain (Ted Pella, Inc.). Excess stain was absorbed with filter paper, and the staining step was repeated on a second drop of Nano-W, which was allowed to sit for one minute before being wicked away and left to dry. Imaging of the stained grids was conducted at 80 kV on a JEOL 1400 transmission electron microscope (JEOL USA Inc.), and images were captured using an AMT XR11 digital camera (AMT).

### Proteomics characterization of OMVs

To confirm that BFT-2 was secreted by ETBF and NTBF+*bft* in OMVs, but not by ETBF Δ*bft* and NTBF OMVs, OMVs were isolated as described above and pelleted and lysed in buffer containing 0.5%SDS, 50mM ammonium bicarbonate, 50mM NaCl, and Halt Protease Inhibitor. Mass spectrometry analysis was performed by the Mass Spectrometry Technology Access Center (MTAC) at Washington University School of Medicine using previously published protocols ^59^. Samples were purified by trichloroacetic acid precipitation. After pelleting and washing with ice-cold acetone, the resulting protein pellet was resuspended in 8M urea and 0.4M ammonium bicarbonate, reduced with 4mM dithiothreitol, and alkylated with 18mM iodoacetamide. The solution was then diluted to <2M urea, and 1µg of trypsin was added for overnight digestion at 37°C. The resulting peptides were desalted using C18 solid-phase extraction spin columns, and eluates were dried under vacuum using a SpeedVac concentrator. Peptides were analyzed by LC–MS/MS using a nanoElute2 coupled to a timsTOF Pro2 mass spectrometer (Bruker Daltonics). Samples were separated on a C18 column (150µm × 25cm, 1.5µm; Bruker). The mobile phases consisted of 0.1% formic acid (FA) in water (mobile phase A) and 0.1%FA in acetonitrile (mobile phase B). Peptides were separated using a 54-min. linear gradient from 3% to 35% mobile phase B, with a total run time of 60 mins. at a constant flow rate of 600nL/min. The timsTOF Pro2 was operated in dia-PASEF mode. Spectra were acquired across an m/z range of 300-1200, using 28 mass windows per cycle. The inverse reduced ion mobility (1/K₀) range was set to 0.65–1.35 V·s/cm². The trapped ion mobility spectrometry (TIMS) analyzer operated at a 100% duty cycle with a ramp time of 75 ms., yielding an overall cycle time of approximately 1.20 seconds. Collision energy was linearly ramped from 59 eV at 1/K₀ = 1.6 to 20 eV at 1/K₀ = 0.6. dia-PASEF data files were analyzed using directDIA™ workflow embedded in Spectronaut 18 (Biognosys AG) against the *Bacteroides fragilis* database. Search parameters included trypsin digestion with cleavage after K or R, allowance for up to two missed cleavages, carbamidomethylation of cysteine (static modification), and variable modifications including oxidized methionine and Protein N-terminal acetylation. Peptide spectrum matches (PSMs), peptides, and protein groups were identified using a false discovery rate (FDR) threshold of 0.01. For quantification, all fragment ions passing the FDR threshold were used, and protein abundances were calculated as the area under the curve of extracted ion chromatograms within the defined peak boundaries for each targeted ion. The mass spectrometry proteomics data have been deposited to the ProteomeXchange Consortium via the PRIDE partner repository^60^ with the dataset identifier PXD073462.

### Colon organoid cell culture

Biopsies from the proximal colon obtained during colonoscopies of three healthy de-identified donors were obtained after informed consent at Dartmouth-Health. Biopsies in PBS (-/-) with EDTA were placed on a shaking rocker in +4°C for 1 hr. to free the crypts from the biopsies. Then the biopsies were vigorously pipetted up and down several times to release crypts. The supernatant containing crypts was collected, and additional PBS was added to the biopsies followed by vigorous pipetting. Crypts in the supernatant were counted, pelleted, and resuspended in 55% Matrigel (Corning Life Sciences) and 45% isolation medium. Cells were added to 24-well plates and then Matrigel solidified cell isolation medium (IntestiCult complete medium with primocin [StemCell Technologies] supplemented with 50ug/mL gentamycin, 50ug/mL vancomycin and 10uM of RhoKi) was added to each well. Cells were maintained in culture until spherical organoids were observed. To develop polarized monolayers of organoids, 1.5 x 10^6^ cells (as spherical organoids) were collected from the Matrigel and seeded on Transwell permeable filters (#3450: Corning Life Sciences for 24-mm Transwell) or 12-mm Snapwell permeable filters (#3801; Corning Life Sciences). Filters were precoated with a 1:20 dilution of Matrigel in cold DPBS for 2 hrs. and then removed just before seeding cells.

IntestiCult media (StemCell Technologies) was added to both apical and basolateral sides of the filters to support cell growth. Once a confluent monolayer was obtained, IntestiCult differentiation media (StemCell Technologies) was used to differentiate the cells prior to experimentation. Cells were differentiated for one week before experiments.

### Exposure of primary human colon organoids to OMVs

Polarized monolayers of colon organoids on 12-mm Snapwell filters were washed with PBS to remove excess mucus, and 1.5 mL of differentiation media was added to the basolateral side. 2 × 10^10^ OMVs from all strains of *B. fragilis*, a concentration of OMVs similar to that measured in conditioned media and in biological fluids ^21,39^, or the same volume of process control (PC) media were applied to the apical side of cells for 1 hr. and then removed. At the end of the 1hr. exposure and then 24 hrs. later the apical and basolateral media were collected for measurements of cytokines [MILLIPLEX MAP Human Cytokine/Chemokine 48-Plex cytokine assay (MilliporeSigma)]. Ussing chamber analysis was used to quantify CFTR Cl^-^ secretion, sodium absorption and TER as described ^61,62^.

### T84 cell culture

T84 cells, a human colon adenocarcinoma cell line (American Type Culture Collection, Manassas, VA) were maintained in DMEM/F-12 media (ATCC) containing l-glutamine supplemented with 5% fetal bovine serum (FBS: Life Technology). For short-circuit current and TER measurements cells were cultured on Transwell filters (Corning, Ithaca, NY). Experiments were performed 21 days postseeding as previously described ^40–42^.

### Ussing chamber measurements of sodium and CFTR Cl^−^ currents

As described in detail elsewhere ^63–66^ cells on Snapwell or Transwell filters were mounted in Ussing chambers. Primary human colon organoids and T84 cells were exposed to either process control (PC: BHIS growth media processed through the OMV isolation procedure, as a control for possible effects on colon organoids due to the media or the OMV isolation process as recommended by the International Society of Extracellular Vesicles ^67^) or OMVs secreted by ETBF, ETBF Δ*bft*, NTBF, or NTBF+*bft* were added to the apical side of monolayers for 1 hr. then CFTR Cl^−^ currents, amiloride-sensitive sodium currents (in colon organoids only since T84 cells do display amiloride-sensitive sodium currents) and TER were measured. Some monolayers were exposed to OMVs for 1 hr., followed by washing OMVs from the apical surface and 24 hrs. later CFTR Cl^−^ currents and TER were measured. To measure TER, a 1 mV pulse was applied to each monolayer and the change in short circuit current was measured and TER was calculated using Ohms Law. Electrogenic sodium currents were reported as amiloride-sensitive currents, since amiloride inhibits the epithelial sodium channel ENaC ^68^. To measure CFTR Cl^-^ secretion, amiloride (50 μM) was first added to the apical solution to inhibit and measure sodium reabsorption. Thereafter, Cl^−^ secretion was stimulated with forskolin (10 μM; Sigma-Aldrich), which increases protein kinase A that phosphorylates and activates CFTR Cl^-^ secretion ^38^.

Finally, thiazolidinone (CFTRinh-172, 20 μM; Millipore, Billerica, MA), an inhibitor of CFTR Cl^−^ channels, was added to the apical solution ^64^. Data are expressed as the forskolin-stimulated, CFTRinh-172 inhibited short circuit current (Isc), which is presented as μA/cm^2^. Data were collected and analyzed using the Data Acquisition Software Acquire and Analyze program (Physiologic Instruments, San Diego, CA).

### Cytokine measurements

Cytokine secretion (apical and basolateral solutions) from primary human colon organoids was measured with the MILLIPLEX MAP Human Cytokine/Chemokine 48-Plex cytokine assay (MilliporeSigma) following the manufactures instructions.

### Immunocytochemistry

Immunocytochemistry was performed to characterize cell types in the colon organoids and to evaluate the effects of OMVs on E-cadherin and claudin-2. Fixation of colon organoids was performed using 4% paraformaldehyde for 15 mins. at room temperature, followed by permeabilization with 0.25% Triton X-100 for 15 mins., blocking using 10% normal goat serum in PBS, and primary antibody incubation at 4°C overnight. After washing, secondary antibodies were added for 1 hr. at room temperature in the dark. SLC26A3 was labeled with a primary anti-mouse antibody (1:250, Santa Cruz #sc-376187), and a secondary goat anti-mouse-488 antibody (1:500, Santa Cruz #sc-376187). CFTR was labeled with a primary anti-mouse antibody (1:100, Millipore # 05-583) and a secondary anti-mouse-488 antibody (1:500). MUC2 was labeled with a primary anti-mouse antibody (1:300), and a secondary anti-mouse-488 antibody (1:500, Abcam ab11197). Chromogranin A was labeled with a primary goat anti rabbit antibody (1:100), and secondary goat anti rabbit-568 antibody (1:1000, Abcam ab15760). Tight junction labeling was performed by fixing cells in 100% cold methanol at -20°C for 5 mins., blocking with 10% goat serum in PBS for 1 hr., and incubating with primary antibodies at 4°C overnight. The secondary antibody was added for 1 hr. at room temperature the following day. Claudin-2 was labeled using a primary goat anti-rabbit antibody (1:100, Thermo Fisher #51-6100), and a secondary goat anti-rabbit-568 antibody (1:500). E-cadherin was labeled using a primary goat anti-mouse antibody (1:250, BD Transduction lab #610182), and a secondary goat anti-mouse-488 antibody (1:500). All antibodies were diluted in PBS with 5% normal goat serum. DAPI staining to label nuclei was performed in fluorshield anti-fade mounting medium. To image the colon organoids, a Yokogawa W1 spinning disk unit coupled to a Nikon Ti-E inverted microscope was used. To examine the effect of OMVs and PC on claudin-2 and E-cadherin, all settings on the confocal microscope were identical for acquiring images for analysis. Analysis of claudin-2 and E-cadherin abundance was performed by an investigator blinded to the treatments. The membrane areas with positive fluorescence signal were segmented using trainable WEKA machine learning plugin in ImageJ ^69^. The segmented objects were thinned to 1 pixel using ImageJ’s “Skeletonize” morphological operation. The mean and median fluorescence intensities were measured from areas defined by the skeleton and widened to 5 pixels using ImageJ’s “Dilate” morphological operation.

### Statistics

Data were analyzed using the R software environment for statistical computing and graphics version 4.1.0 ^70^, Ingenuity Pathway Analysis and Prism (GraphPad). Statistical significance was calculated using either a mixed effect linear model, or likelihood ratio tests on gene-wise negative binomial generalized linear models, as indicated in the figure legends. Data were visualized, and figures were created using Prism or ggplot.

## Author contributions

PS, RB, AN, BAS, CR, TTH and BR designed the research studies. PS, RB, AN, CR, TTH and ZS conducted experiments on human colon organoids and T84 cells, acquired data, and analyzed data. TBG recruited human subjects, provided informed consent and collected clinical biopsies. YAG and BKC performed the proteomics analysis of OMVs. Electron microscopy of OMVs was performed by DJT and Brad Vietje at the Microscopy Imaging Center at the University of Vermont (RRID# SCR_018821). BAS wrote the first and revised draft of the manuscript. All authors contributed to the article and approved the submitted version.

## IRB statement

The Dartmouth Committee for the Protection of Human Subjects has determined that using colon biopsies in this study is not considered human subject research because they contain no patient identifiers. All tissues used for isolation were obtained under informed consent and conform to HIPAA regulations to protect the privacy of the donor’s personally identifiable information.

## Data availability

The mass spectrometry proteomics data are available in the Supplemental Tables and have been deposited to the ProteomeXchange Consortium via the PRIDE partner repository^60^ with the dataset identifier PXD073462. All other reasonable requests for additional data are available from the corresponding author.

## Acknowledgements

Cynthia Sears, M.D. (Johns Hopkins University) generously provided the ETBF Δ*bft* strain. Scott Palisoul (Dartmouth Pathology Shared Resources) created frozen cross sections of 2D colon organoids. We thank Brad Vietje for the electron micrograph’s and Katja Koeppen, Ph.D. for analysis of the T84 data.

## SUPPLEMENTAL TABLES

**Supplemental Table 1.** Mass spectrometry quantification of proteins detected exclusively in *B. fragilis* NTBF OMVs.

**Supplemental Table 2.** Mass spectrometry quantification of proteins detected exclusively in *B. fragilis* ETBF OMVs.

**Supplemental Table 3.** Mass spectrometry quantification of proteins detected in both *B. fragilis* NTBF and *B. fragilis* ETBF OMVs.

**Supplemental Table 4.** Differential expression results of proteins identified in mass spectrometry for both *B. fragilis* ETBF and *B. fragilis* NTBF

## References

1 Jasemi, S. et al. Biological Mechanisms of Enterotoxigenic Bacteroides fragilis Toxin: Linking Inflammation, Colorectal Cancer, and Clinical Implications. Toxins (Basel) 17 (2025). 10.3390/toxins17060305

2 Ferranti, E. P., Dunbar, S. B., Dunlop, A. L. & Corwin, E. J. 20 things you didn’t know about the human gut microbiome. J Cardiovasc Nurs 29, 479–481 (2014). 10.1097/JCN.0000000000000166

3 Valguarnera, E. & Wardenburg, J. B. Good Gone Bad: One Toxin Away From Disease for Bacteroides fragilis. J Mol Biol 432, 765–785 (2020). 10.1016/j.jmb.2019.12.003

4 Human Microbiome Project, C. Structure, function and diversity of the healthy human microbiome. Nature 486, 207–214 (2012). 10.1038/nature11234

5 Young, V. B. The role of the microbiome in human health and disease: an introduction for clinicians. BMJ 356, j831 (2017). 10.1136/bmj.j831

6 Saha, K., Zhou, Y. & Turner, J. R. Tight junction regulation, intestinal permeability, and mucosal immunity in gastrointestinal health and disease. Curr Opin Gastroenterol 41, 46–53 (2025). 10.1097/MOG.0000000000001066

7 Hill, C. A. et al. Bacteroides fragilis toxin expression enables lamina propria niche acquisition in the developing mouse gut. Nat Microbiol 9, 85–94 (2024). 10.1038/s41564-023-01559-9

8 Sears, C. L. et al. Association of enterotoxigenic Bacteroides fragilis infection with inflammatory diarrhea. Clin Infect Dis 47, 797–803 (2008). 10.1086/591130

9 Kim, W. S. et al. Bacteroides fragilis Toxin Induces Sequential Proteolysis of E-Cadherin and Inflammatory Response in Mouse Intestinal Epithelial Cell Line. Microorganisms 13 (2025). 10.3390/microorganisms13040781

10 Hwang, S., Gwon, S. Y., Kim, M. S., Lee, S. & Rhee, K. J. Bacteroides fragilis Toxin Induces IL-8 Secretion in HT29/C1 Cells through Disruption of E-cadherin Junctions. Immune Netw 13, 213–217 (2013). 10.4110/in.2013.13.5.213

11 Guo, Y. et al. Mechanistic diversity of Bacteroides fragilis toxins and neutralization with single domain antibody. Cell Chem Biol 33, 102–116 e106 (2026). 10.1016/j.chembiol.2025.12.009

12 Zakharzhevskaya, N. B. et al. Interaction of Bacteroides fragilis Toxin with Outer Membrane Vesicles Reveals New Mechanism of Its Secretion and Delivery. Front Cell Infect Microbiol 7, 2 (2017). 10.3389/fcimb.2017.00002

13 Rong, Y. et al. Partially differentiated ileal and distal-colonic human F508del-cystic fibrosis-enteroids secrete fluid in response to forskolin and linaclotide. iScience 28, 112453 (2025). 10.1016/j.isci.2025.112453

14 Negussie, A. B., Dell, A. C., Davis, B. A. & Geibel, J. P. Colonic Fluid and Electrolyte Transport 2022: An Update. Cells 11 (2022). 10.3390/cells11101712

15 Riegler, M. et al. Bacteroides fragilis toxin 2 damages human colonic mucosa in vitro. Gut 44, 504–510 (1999). 10.1136/gut.44.4.504

16 Obiso, R. J., Jr., Azghani, A. O. & Wilkins, T. D. The Bacteroides fragilis toxin fragilysin disrupts the paracellular barrier of epithelial cells. Infect Immun 65, 1431–1439 (1997). 10.1128/iai.65.4.1431-1439.1997

17 Wu, S., Rhee, K. J., Zhang, M., Franco, A. & Sears, C. L. Bacteroides fragilis toxin stimulates intestinal epithelial cell shedding and gamma-secretase-dependent E-cadherin cleavage. J Cell Sci 120, 1944–1952 (2007). 10.1242/jcs.03455

18 Sheikh, A. et al. Outer membrane vesicles from Bacteroides fragilis contain coding and non-coding small RNA species that modulate inflammatory signaling in intestinal epithelial cells. bioRxiv (2025). 10.1101/2025.06.25.661399

19 Kharlampieva, D. D. et al. Purification and characterisation of recombinant Bacteroides fragilis toxin-2. Biochimie 95, 2123–2131 (2013). 10.1016/j.biochi.2013.08.005

20 Chambers, F. G. et al. Bacteroides fragilis toxin exhibits polar activity on monolayers of human intestinal epithelial cells (T84 cells) in vitro. Infect Immun 65, 3561–3570 (1997). 10.1128/iai.65.9.3561-3570.1997

21 Charpentier, L. A. et al. Bacterial Outer Membrane Vesicles and Immune Modulation of the Host. Membranes (Basel) 13 (2023). 10.3390/membranes13090752

22 Li, Z. et al. P. aeruginosa tRNA-fMet halves secreted in outer membrane vesicles suppress lung inflammation in cystic fibrosis. Am J Physiol Lung Cell Mol Physiol 326, L574–L588 (2024). 10.1152/ajplung.00018.2024

23 Koeppen, K. et al. A Novel Mechanism of Host-Pathogen Interaction through sRNA in Bacterial Outer Membrane Vesicles. PLoS Pathog 12, e1005672 (2016). 10.1371/journal.ppat.1005672

24 Yang, Y. et al. Bacteroides Fragilis-Derived Outer Membrane Vesicles Deliver MiR-5119 and Alleviate Colitis by Targeting PD-L1 to Inhibit GSDMD-Mediated Neutrophil Extracellular Trap Formation. Adv Sci (Weinh), e00781 (2025). 10.1002/advs.202500781

25 Shagaleeva, O. Y. et al. Bacteroides vesicles promote functional alterations in the gut microbiota composition. Microbiol Spectr 12, e0063624 (2024). 10.1128/spectrum.00636-24

26 Patrick, S., McKenna, J. P., O’Hagan, S. & Dermott, E. A comparison of the haemagglutinating and enzymic activities of Bacteroides fragilis whole cells and outer membrane vesicles. Microb Pathog 20, 191–202 (1996). 10.1006/mpat.1996.0018

27 Sharpe, S. W., Kuehn, M. J. & Mason, K. M. Elicitation of epithelial cell-derived immune effectors by outer membrane vesicles of nontypeable Haemophilus influenzae. Infect Immun 79, 4361–4369 (2011). 10.1128/IAI.05332-11

28 Round, J. L. et al. The Toll-like receptor 2 pathway establishes colonization by a commensal of the human microbiota. Science 332, 974–977 (2011). 10.1126/science.1206095

29 Lee, S. M. et al. Bacterial colonization factors control specificity and stability of the gut microbiota. Nature 501, 426–429 (2013). 10.1038/nature12447

30 Sears, C. L., Geis, A. L. & Housseau, F. Bacteroides fragilis subverts mucosal biology: from symbiont to colon carcinogenesis. J Clin Invest 124, 4166–4172 (2014). 10.1172/JCI72334

31 Taglieri, M. et al. Colorectal Organoids: Models, Imaging, Omics, Therapy, Immunology, and Ethics. Cells 14 (2025). 10.3390/cells14060457

32 Sen, A. et al. Comprehensive gene expression analysis of organoid-derived healthy human colonic epithelium and cancer cell line stimulated with live probiotic bacteria. Sci Rep 15, 22325 (2025). 10.1038/s41598-025-07391-x

33 Rikkert, L. G., Nieuwland, R., Terstappen, L. & Coumans, F. A. W. Quality of extracellular vesicle images by transmission electron microscopy is operator and protocol dependent. J Extracell Vesicles 8, 1555419 (2019). 10.1080/20013078.2018.1555419

34 Noble, J. M. et al. Direct comparison of optical and electron microscopy methods for structural characterization of extracellular vesicles. J Struct Biol 210, 107474 (2020). 10.1016/j.jsb.2020.107474

35 Weiss, A. et al. Transmission Electron Microscopy-based characterization of Extracellular Vesicles from plasma and serum from Parkinson s Disease patients. Cell Commun Signal 23, 395 (2025). 10.1186/s12964-025-02383-w

36 Zakharzhevskaya, N. B. et al. Outer membrane vesicles secreted by pathogenic and nonpathogenic Bacteroides fragilis represent different metabolic activities. Sci Rep 7, 5008 (2017). 10.1038/s41598-017-05264-6

37 Parikh, K. et al. Colonic epithelial cell diversity in health and inflammatory bowel disease. Nature 567, 49–55 (2019). 10.1038/s41586-019-0992-y

38 Ghigo, A., De Santi, C., Hart, M., Mitash, N. & Swiatecka-Urban, A. Cell signaling and regulation of CFTR expression in cystic fibrosis cells in the era of high efficiency modulator therapy. J Cyst Fibros 22 **Suppl 1**, S12–S16 (2023). 10.1016/j.jcf.2022.12.015

39 Stanton, B. A. Extracellular Vesicles and Host-Pathogen Interactions: A Review of Inter-Kingdom Signaling by Small Noncoding RNA. Genes (Basel) 12 (2021). 10.3390/genes12071010

40 Howe, K. L., Wang, A., Hunter, M. M., Stanton, B. A. & McKay, D. M. TGFbeta down-regulation of the CFTR: a means to limit epithelial chloride secretion. Exp Cell Res 298, 473–484 (2004). 10.1016/j.yexcr.2004.04.026

41 Maitra, R., Shaw, C. M., Stanton, B. A. & Hamilton, J. W. Increased functional cell surface expression of CFTR and DeltaF508-CFTR by the anthracycline doxorubicin. Am J Physiol Cell Physiol 280, C1031–1037 (2001). 10.1152/ajpcell.2001.280.5.C1031

42 Maitra, R., Shaw, C. M., Stanton, B. A. & Hamilton, J. W. Functional enhancement of CFTR expression by mitomycin C. Cell Physiol Biochem 11, 93–98 (2001). 10.1159/000047796

43 Pierce, J. V., Fellows, J. D., Anderson, D. E. & Bernstein, H. D. A clostripain-like protease plays a major role in generating the secretome of enterotoxigenic Bacteroides fragilis. Mol Microbiol 115, 290–304 (2021). 10.1111/mmi.14616

44 Choi, V. M. et al. Activation of Bacteroides fragilis toxin by a novel bacterial protease contributes to anaerobic sepsis in mice. Nat Med 22, 563–567 (2016). 10.1038/nm.4077

45 Elhenawy, W., Debelyy, M. O. & Feldman, M. F. Preferential packing of acidic glycosidases and proteases into Bacteroides outer membrane vesicles. MBio 5, e00909–00914 (2014). 10.1128/mBio.00909-14

46 France, M. M. & Turner, J. R. The mucosal barrier at a glance. J Cell Sci 130, 307–314 (2017). 10.1242/jcs.193482

47 Citi, S. et al. A short guide to the tight junction. J Cell Sci 137 (2024). 10.1242/jcs.261776

48 Han, Y. W. et al. Interactions between periodontal bacteria and human oral epithelial cells: Fusobacterium nucleatum adheres to and invades epithelial cells. Infect Immun 68, 3140–3146 (2000). 10.1128/IAI.68.6.3140-3146.2000

49 Antosca, K. M. et al. Altered Stool Microbiota of Infants with Cystic Fibrosis Shows a Reduction in Genera Associated with Immune Programming from Birth. J Bacteriol 201 (2019). 10.1128/JB.00274-19

50 Lee, C. G. et al. Bacteroides fragilis Toxin Induces Intestinal Epithelial Cell Secretion of Interleukin-8 by the E-Cadherin/beta-Catenin/NF-kappaB Dependent Pathway. Biomedicines 10 (2022). 10.3390/biomedicines10040827

51 Choi, J. W., Kim, S. C., Hong, S. H. & Lee, H. J. Secretable Small RNAs via Outer Membrane Vesicles in Periodontal Pathogens. J Dent Res 96, 458–466 (2017). 10.1177/0022034516685071

52 Franco, A. A. The Bacteroides fragilis pathogenicity island is contained in a putative novel conjugative transposon. J Bacteriol 186, 6077–6092 (2004). 10.1128/JB.186.18.6077-6092.2004

53 Wu, S. et al. A human colonic commensal promotes colon tumorigenesis via activation of T helper type 17 T cell responses. Nat Med 15, 1016–1022 (2009). 10.1038/nm.2015

54 Chan, J. L. et al. Non-toxigenic Bacteroides fragilis (NTBF) administration reduces bacteria-driven chronic colitis and tumor development independent of polysaccharide A. Mucosal Immunol 12, 164–177 (2019). 10.1038/s41385-018-0085-5

55 Chung, L. et al. Bacteroides fragilis Toxin Coordinates a Pro-carcinogenic Inflammatory Cascade via Targeting of Colonic Epithelial Cells. Cell Host Microbe 23, 203–214 e205 (2018). 10.1016/j.chom.2018.01.007

56 Lim, B., Zimmermann, M., Barry, N. A. & Goodman, A. L. Engineered Regulatory Systems Modulate Gene Expression of Human Commensals in the Gut. Cell 169, 547–558 e515 (2017). 10.1016/j.cell.2017.03.045

57 Degnan, P. H., Barry, N. A., Mok, K. C., Taga, M. E. & Goodman, A. L. Human gut microbes use multiple transporters to distinguish vitamin B(1)(2) analogs and compete in the gut. Cell Host Microbe 15, 47–57 (2014). 10.1016/j.chom.2013.12.007

58 Bauman, S. J. & Kuehn, M. J. Purification of outer membrane vesicles from Pseudomonas aeruginosa and their activation of an IL-8 response. Microbes Infect 8, 2400–2408 (2006). 10.1016/j.micinf.2006.05.001

59 Koeppen, K. et al. An rRNA fragment in extracellular vesicles secreted by human airway epithelial cells increases the fluoroquinolone sensitivity of P. aeruginosa. Am J Physiol Lung Cell Mol Physiol 325, L54–L65 (2023). 10.1152/ajplung.00150.2022

60 Perez-Riverol, Y. et al. The PRIDE database at 20 years: 2025 update. Nucleic Acids Res 53, D543–D553 (2025). 10.1093/nar/gkae1011

61 Hampton, T. H., Koeppen, K., Bashor, L. & Stanton, B. A. Selection of reference genes for quantitative PCR: identifying reference genes for airway epithelial cells exposed to Pseudomonas aeruginosa. Am J Physiol Lung Cell Mol Physiol 319, L256–L265 (2020). 10.1152/ajplung.00158.2020

62 Hampton, T. H. et al. Gene expression responses of CF airway epithelial cells exposed to elexacaftor/tezacaftor/ivacaftor (ETI) suggest benefits beyond improved CFTR channel function. Am J Physiol Lung Cell Mol Physiol. Dec 1;327(6):L905–L916. (2024). doi: 10.1152/ajplung.00272.2024.

63 Stanton, B. A., Coutermarsh, B., Barnaby, R. & Hogan, D. Pseudomonas aeruginosa Reduces VX-809 Stimulated F508del-CFTR Chloride Secretion by Airway Epithelial Cells. PLoS One 10, e0127742 (2015). 10.1371/journal.pone.0127742

64 Barnaby, R., Koeppen, K. & Stanton, B. A. Cyclodextrins reduce the ability of Pseudomonas aeruginosa outer-membrane vesicles to reduce CFTR Cl(-) secretion. Am J Physiol Lung Cell Mol Physiol 316, L206–L215 (2019). 10.1152/ajplung.00316.2018

65 Talebian, L., Coutermarsh, B., Channon, J. Y. & Stanton, B. A. Corr4A and VRT325 do not reduce the inflammatory response to P. aeruginosa in human cystic fibrosis airway epithelial cells. Cell Physiol Biochem 23, 199–204 (2009). 10.1159/000204108

66 Stanton, B. A. Effects of Pseudomonas aeruginosa on CFTR chloride secretion and the host immune response. Am J Physiol Cell Physiol 312, C357–C366 (2017). 10.1152/ajpcell.00373.2016

67 Welsh, J. A. et al. Minimal information for studies of extracellular vesicles (MISEV2023): From basic to advanced approaches. J Extracell Vesicles 13, e12404 (2024). 10.1002/jev2.12404

68 Berdiev, B. K., Shlyonsky, V. G., Karlson, K. H., Stanton, B. A. & Ismailov, II. Gating of amiloride-sensitive Na(+) channels: subunit-subunit interactions and inhibition by the cystic fibrosis transmembrane conductance regulator. Biophys J 78, 1881–1894 (2000). 10.1016/S0006-3495(00)76737-3

69 Arganda-Carreras, I. et al. Trainable Weka Segmentation: a machine learning tool for microscopy pixel classification. Bioinformatics 33, 2424–2426 (2017). 10.1093/bioinformatics/btx180

70 R Core Team (2017) R: A Language and Environment for Statistical Computing. https://www.R-project.org/

